# Tyrosine phosphorylation and dimerization cooperatively activate NAMPT to enable NAD+ synthesis in cancer

**DOI:** 10.64898/2026.08.13.744642

**Authors:** Johnvesly Basappa, Cosimo Lobello, Anneliese M Faustino, Cristina Uribe-Alvarez, David Rushmore, Niel Sen, Li Wang, Andrey Efimov, Kathy Q. Cai, Jaime L Schneider, Lori Rink, Aaron N Hata, Luca Mologni, Wujuan Zhang, Aaron R. Goldman, Hsin-Yao Tang, Reza Nejati, Roland Dunbrack, Jonathan Chernoff, Joseph A Baur, Mariusz A. Wasik

## Abstract

Nicotinamide phosphoribosyltransferase (NAMPT), the rate-limiting enzyme in the NAD⁺ salvage pathway, is frequently upregulated in cancer, yet mechanisms regulating its catalytic activity remain undefined. We identify NAMPT as a direct substrate of multiple proto-oncogenic tyrosine kinases, including ALK, insulin receptor, IGF1R, and PDGFRA. Phosphoproteomics identified NAMPT Y188 as the major phosphorylation site, including the oncogenic fusion kinase NPM1::ALK. NAMPT interacted with NPM1::ALK in the cytoplasm, nucleus, and mitochondria, while Y188 phosphorylation enhanced catalytic activity, NMN/NAD⁺ biosynthesis, and downstream metabolism. Conversely, the Y188F mutant reduced enzymatic activity, proliferation, and clonogenicity, whereas disrupting dimerization similarly impaired phosphorylation and function. Interactome analyses showed phosphorylation and dimerization cooperatively remodel NAMPT-associated networks, enriching phosphorylated dimers for metabolic/redox regulators and monomeric NAMPT for ribosome biogenesis. NAMPT inhibition suppressed the growth of both ALK inhibitor-sensitive and -resistant lymphoma cells and enhanced the efficacy of ALK inhibition, revealing kinase-dependent NAMPT activation as a metabolic vulnerability in oncogene-driven cancers.

## INTRODUCTION

Tyrosine kinases are central regulators of oncogenic signaling that control cellular behavior through post-translational modifications. Although tyrosine phosphorylation accounts for less than 1% of all phosphorylation sites, it exerts disproportionate effects on protein function, cellular signaling and cancer progression by modulating enzymatic activity, protein-protein interactions and downstream signaling networks (*1–5*). Beyond their established roles in proliferation and survival, tyrosine kinases have emerged as key regulators of cancer metabolism. Accumulating evidence, including our own work, has demonstrated that tyrosine kinase-mediated phosphorylation of metabolic enzymes, including ATP citrate lyase (ACLY) (*6, 7*), guanylate kinase 1 (GUK1) (*8*), Pyruvate Kinase M2 (PKM2)(*9*), Pyruvate Dehydrogenase Kinase (PDK1)(*10*) and Lactate Dehydrogenase (LDHA)(*11*), promotes metabolic reprogramming in cancer cells. These modifications alter enzymatic activity, substrate utilization and pathway integration to support the anabolic and bioenergetic demands of tumor growth. In addition, tyrosine phosphorylation can create docking sites for SH2- and PTB-domain-containing proteins, thereby coupling metabolic regulation to broader signaling networks (*12, 13*). Anaplastic lymphoma kinase (ALK) is a receptor tyrosine kinase with well-established oncogenic functions. ALK was originally identified in anaplastic large cell lymphoma (ALCL) through its fusion with nucleophosmin (NPM), generating the constitutively active NPM1::ALK oncoprotein (*14*). Subsequently, ALK rearrangements have been identified in multiple malignancies, including non-small cell lung cancer (NSCLC) through EML4–ALK fusions (*15*), inflammatory myofibroblastic tumors (*16*) and neuroblastoma, where activating mutations of full-length ALK drive tumorigenesis (*17*). Together with leukocyte tyrosine kinase (LTK) and ROS1, ALK belongs to the insulin receptor superfamily of receptor tyrosine kinases(*18*). ALCL is an aggressive subtype of non-Hodgkin lymphoma, accounting for approximately 3% of adult and 10–15% of pediatric cases (*19*). The potent oncogenic activity of NPM1::ALK has been demonstrated in both cell-based and transgenic animal models, in which its expression is sufficient to induce malignant transformation and tumor formation (*20, 21*).

Nicotinamide phosphoribosyltransferase (NAMPT) is the rate-limiting enzyme of the NAD+ salvage pathway, catalyzing the conversion of nicotinamide (NAM) to nicotinamide mononucleotide (NMN), which is subsequently converted to NAD+ by nicotinamide mononucleotide adenylyltransferases (NMNATs) (*22, 23*). NAD+ biosynthesis occurs through the de novo, Preiss–Handler and salvage pathways, with the salvage pathway serving as the predominant source of NAD+ in most mammalian tissues(*24*). Because nicotinamide is the principal precursor for NAD+ synthesis in higher vertebrates, NAMPT occupies a central position in maintaining cellular NAD+ homeostasis (*25, 26*). Consistent with this role, our previous studies demonstrated that NPM1::ALK upregulates in ALCL expression of *NAMPT* gene, implicating NAMPT in ALK-driven oncogenesis (*27*). Human NAMPT is a 55-kDa protein that functions primarily as a homodimer, with the active site formed at the dimer interface (*28*). In contrast, extracellular NAMPT predominantly exists as a monomer and is reported to function as a cytokine-like signaling molecule independent of NAD+ biosynthesis(*28*), but has also been observed in adipose-derived extracellular vesicles that target the hypothalamus and correlated with NAD+ in that tissue (*29*). NAMPT over expression has been reported across numerous solid tumors, including prostate, glioma, melanoma, lung, colon, breast, thyroid, kidney, and pancreatic cancer as well as hematological malignancies, including BTK-driven cancers(*30*), chronic lymphocytic leukemia (*31*, *32*). multiple myeloma (*33*), acute myeloid leukemia (*34–36*), chronic myeloid leukemia (*37*), T-cell leukemia/lymphoma (*38*), diffuse large B-cell lymphoma, follicular lymphoma and mantle cell lymphoma (*39*).

Despite the importance of NAMPT in cancer metabolism, the mechanisms by which post-translational modifications regulate its enzymatic activity remain largely undefined. Moreover, the roles of its monomeric and dimeric states, as well as its protein–protein interaction network, are poorly understood. Here, we identify NAMPT as a direct substrate of oncogenic tyrosine kinases and investigate how tyrosine phosphorylation influences NAMPT dimerization, enzymatic activity, NAD⁺ biosynthesis, and protein–protein interactions using NAMPT immunoprecipitation coupled with LC–MS/MS-based metabolomic and proteomic analysis.

## RESULTS

### Oncogenic tyrosine kinases directly phosphorylate NAMPT on multiple tyrosine residues

We hypothesized that oncogenic tyrosine kinases directly phosphorylate NAMPT to enhance its activity and promote metabolic reprogramming in hematological malignancies (Fig. 1A). To test this, we performed in vitro kinase assays using recombinant human NAMPT and a panel of active tyrosine kinases, including ALK, BTK, SYK, SRC, FYN, LCK, FLT3, ABL1, and EGFR, with ERK1 and AKT serving as serine/threonine kinase controls (Extended Data Fig. 1B). Immunoblotting with anti-phosphotyrosine (pY100) confirmed robust tyrosine phosphorylation of NAMPT by multiple tyrosine kinases, whereas ERK1 and AKT failed to phosphorylate NAMPT, demonstrating that NAMPT is a direct substrate of oncogenic tyrosine kinases (Fig. 1B). To identify phosphorylation sites, recombinant NAMPT was incubated with SYK or BTK, two kinases frequently activated in hematologic malignancies (Fig. 1C). Efficient phosphorylation was confirmed by immunoblotting prior to LC-MS/MS phosphoproteomic analysis (Fig. 1D). We identified ten tyrosine phosphorylation sites (Y34, Y54, Y87, Y103, Y108, Y188, Y195, Y341, Y453, and Y471), including five previously unreported sites (Y87, Y103, Y108, Y341, and Y471) (Fig. 1E). Quantitative analysis revealed distinct phosphorylation profiles following SYK- and BTK-mediated phosphorylation (Fig. 1F and G), while mapping of phosphosites across the NAMPT sequence indicated widespread kinase-dependent regulation (Fig. 1H). Among the identified residues, Y188 emerged as a priority candidate because it was robustly phosphorylated in our in vitro assays and identified as such in public phosphoproteomic database (https://phosphosite.org). However, the upstream kinase responsible for this modification and its functional significance have remained unknown. Here, we demonstrate that ALK directly phosphorylates Y188, enhancing NAMPT enzymatic activity and promoting metabolic signaling in lymphoma. Structural analysis of the NAMPT dimer (PDB: 2E5B) revealed that residue Y188 is positioned on the protein surface in a region that is accessible for potential post-translational modification. The docking results showed that several residues are located within 10 Å of Y188, including K216, which is situated near the tyrosine residue. Given its positively charged side chain, K216 may establish a favorable electrostatic interaction with phosphorylated Y188, potentially stabilizing the phosphorylated state. This spatial arrangement suggests that phosphorylation at Y188 could influence the local interaction network and may contribute to modulation of NAMPT structure or function (Fig. 1I). Sequence alignment demonstrated that Y188 is highly conserved across vertebrates, supporting its functional importance (Fig. 1J). To validate Y188 phosphorylation in cells, HEK293T cells expressing HA-tagged NAMPT-WT or the phospho-deficient Y188F mutant were transfected with SYK, SRC, or BTK and treated with the corresponding kinase inhibitors. Immunoprecipitation followed by phospho-Y188 immunoblotting showed that expression of each kinase increased Y188 phosphorylation, whereas entospletinib, dasatinib, or ibrutinib markedly reduced phosphorylation. The Y188F mutant abolished phospho-Y188 detection, confirming antibody specificity and validating Y188 as a bona fide phosphorylation site (Fig. 1K). Together, these findings establish NAMPT as a direct substrate of multiple oncogenic tyrosine kinases and identify Y188 as an evolutionarily conserved regulatory phosphorylation site linking tyrosine kinase signaling to NAD metabolism.

**Fig. 1.**
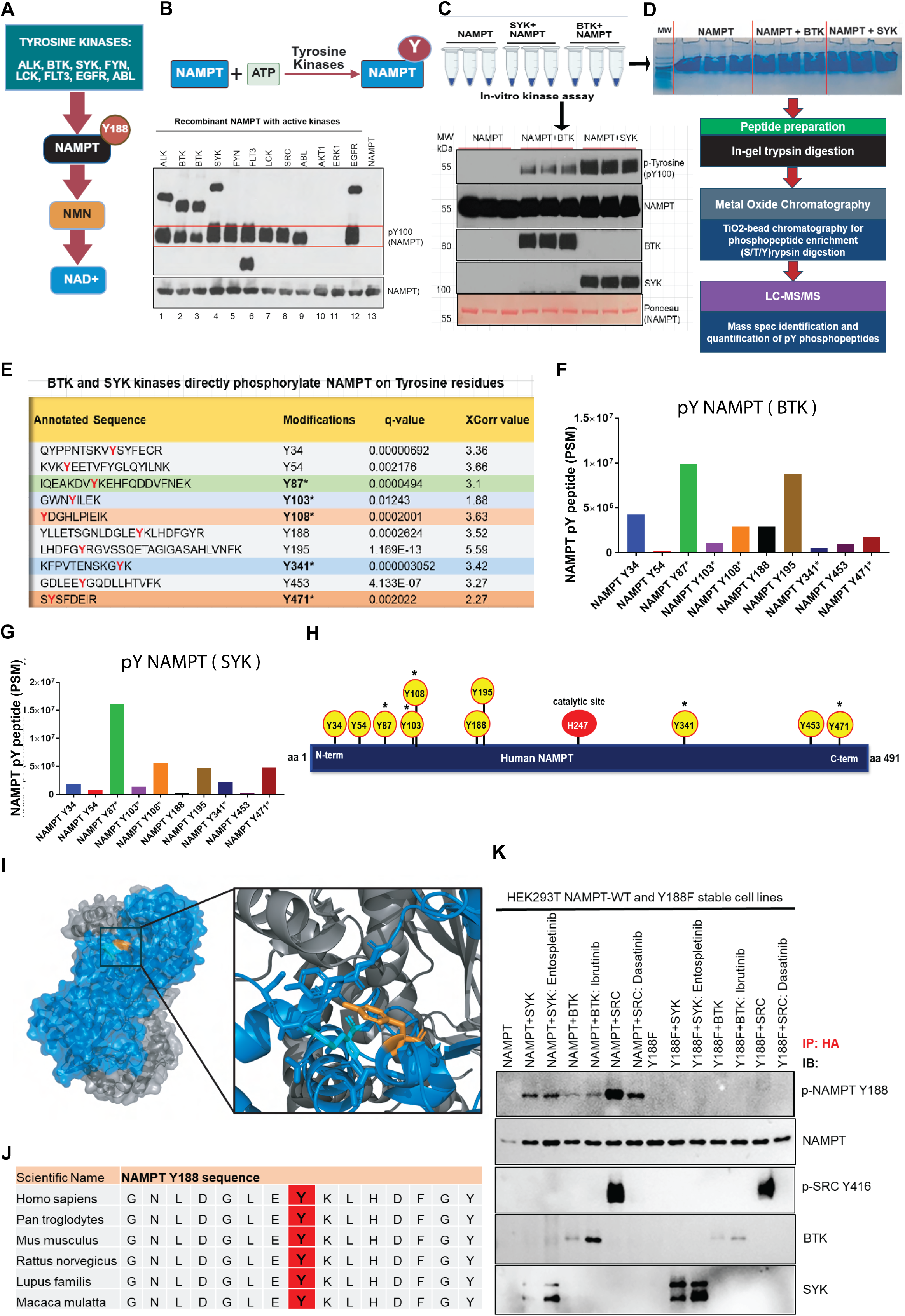
| Oncogenic tyrosine kinases directly phosphorylate NAMPT on multiple tyrosine residues and identify Y188 as a conserved regulatory site. **(A)** Schematic illustrating the hypothesis that oncogenic tyrosine kinases directly phosphorylate NAMPT. **(B)** Schematic of in-vitro kinase assays using recombinant human NAMPT incubated with the indicated active tyrosine kinases (ALK, BTK, SYK, SRC, FYN, LCK, FLT3, ABL1 and EGFR) or the serine/threonine kinases ERK1 and AKT as controls. Tyrosine phosphorylation of NAMPT was detected by immunoblotting with anti-phosphotyrosine (pY100) antibody. **(C)** Large scale in-vitro kinase assay using SYK and BTK kinase and Immunoblot validation of efficient NAMPT phosphorylation by SYK and BTK before LC–MS/MS analysis. **(D)** Experimental workflow for phosphosite identification. Recombinant NAMPT was phosphorylated in vitro by SYK or BTK prior to phosphoproteomic analysis by LC–MS/MS. **(E)**, Identification of ten tyrosine phosphorylation sites on NAMPT by LC–MS/MS and newly identified phosphosites are indicated with *. **(F and G)** Quantitative phosphoproteomic analysis showing relative phosphorylation abundance of individual NAMPT tyrosine residues following phosphorylation by SYK (F) or BTK (F). **(H)** Distribution of identified tyrosine phosphorylation sites across the NAMPT protein sequence, illustrating kinase-dependent phosphorylation at multiple regions of the enzyme. **(I)** Structure of the NAMPT dimer (PDB: 2E5B) with monomers colored blue and gray and Y188 is highlighted in orange. The inset provides a zoomed view of the region surrounding Y188, with residues located within 10 Å displayed with sticks visible. K216, shown in cyan, is positioned in proximity to Y188. **(J)** Multiple sequence alignment of NAMPT orthologues demonstrating evolutionary conservation of Y188 across vertebrate species. **(K)** Cellular validation of Y188 phosphorylation. HEK293T cells expressing HA-tagged NAMPT-WT or the phospho-deficient NAMPT-Y188F mutant together with SYK, SRC or BTK were treated with the corresponding kinase inhibitors (entospletinib, dasatinib or ibrutinib). HA-immunoprecipitates were analyzed by immunoblotting using a phospho-Y188-specific antibody. Kinase expression increased Y188 phosphorylation, whereas kinase inhibition reduced phosphorylation. The Y188F mutation abolished phospho-Y188 signal, confirming antibody specificity and validating Y188 as a bona fide tyrosine phosphorylation site.

### NPM1::ALK phosphorylates NAMPT at the highly-conserved Y188 residue

A schematic illustrating NPM1::ALK-mediated phosphorylation of NAMPT Y188 in the presence or absence of ALK inhibition is shown (Fig. 2A and B). To investigate Y188 phosphorylation, HEK293T cells stably expressing HA-tagged wild-type (dimeric) or dimerization-defective SS-AA (monomeric) NAMPT together with constitutively active NPM1::ALK were analyzed (Fig. 2C). Immunoblotting revealed robust Y188 phosphorylation in wild-type NAMPT, whereas phosphorylation was markedly reduced in the SS-AA mutant, indicating that NAMPT dimerization promotes efficient phosphorylation by NPM1::ALK (Fig. 2D). Densitometric analysis confirmed significantly decreased p-NAMPT Y188 levels in monomeric NAMPT (Fig. 2E). To comprehensively identify NPM1::ALK-dependent phosphorylation sites, HA immunoprecipitates from wild-type and SS-AA NAMPT-expressing cells were subjected to LC–MS/MS phosphoproteomic analysis. Four phosphotyrosine residues (Y34, Y175, Y188, and Y403) were identified, with Y188 representing the dominant phosphosite, exhibiting substantially greater phosphopeptide abundance than Y34, Y175, or Y403 (Fig. 2F). Representative MS/MS spectra confirmed confident identification of all four phosphotyrosine-containing peptides (Fig. 2G). The schematic mapping of the identified phosphosites onto dimeric and monomeric NAMPT models showed that Y188 is positioned adjacent to the dimer interface, supporting a potential role for phosphorylation in regulating NAMPT conformation and activity (Fig. 2H and I). High-confidence of phosphorylation at Y188 was confirmed by LC–MS/MS fragmentation spectra (Fig. 2J). In parallel, phosphorylation of NPM1::ALK was quantified to assess whether NAMPT conformation influences ALK signaling. Multiple ALK autophosphorylation sites (Y1078, Y1096, Y1131, Y1239, Y1358, Y1507, and Y1584) were detected, all of which were consistently reduced in cells expressing monomeric SS-AA NAMPT (Fig. 2K). Together, these findings identify NAMPT as a direct substrate of NPM1::ALK, establish Y188 as the predominant phosphorylation site, and reveal reciprocal regulation between NAMPT dimerization and ALK kinase signaling.

**Fig. 2.**
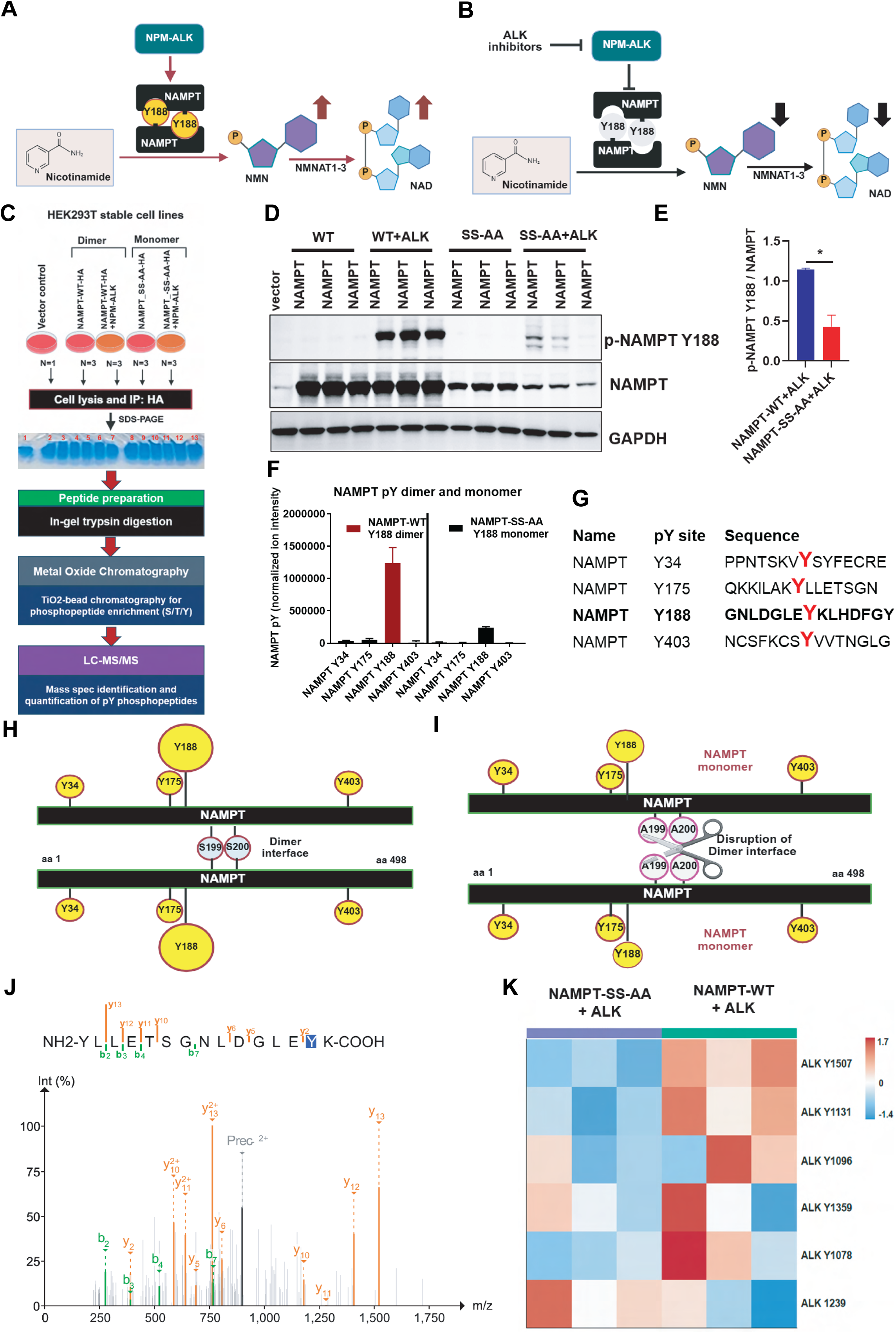
NPM1::ALK phosphorylates NAMPT at the conserved Y188 residue. **(A and B)** Schematic illustrating NPM–ALK-mediated phosphorylation of NAMPT at Y188 and the experimental design used to evaluate Y188 phosphorylation in the presence or absence of ALK inhibition. **(C)** Experimental workflow for phosphoproteomic analysis of HA-tagged wild-type (WT) NAMPT and dimerization-defective SS-AA NAMPT expressed in HEK293T cells together with constitutively active NPM–ALK. (**D**) Immunoblot analysis of phospho-NAMPT Y188 in HA-NAMPT-WT and HA-NAMPT-SS-AA cells expressing NPM1::ALK. Lysates were probed with the indicated antibodies. Dimeric NAMPT exhibits substantially higher Y188 phosphorylation than the monomeric SS-AA mutant. **(E)** Densitometric quantification of p-NAMPT Y188 (dimer)/NAMPT and monomer from figure d. (**F**) Quantification of phosphotyrosine-containing NAMPT peptides identified by LC–MS/MS. Y188 represents the most abundant phosphotyrosine site. Four phosphotyrosine sites were detected: Y34, Y175, Y188, and Y403. (**G)** Representative MS/MS spectra of phosphopeptides corresponding to NAMPT Y34, Y175, Y188, and Y403. (**H and I**) Schematic mapping of identified phosphotyrosine residues onto dimeric **(H)** and monomeric **(I)** NAMPT models. Y188 is located proximal to the NAMPT dimerization interface. (**J)** Representative LC–MS/MS fragmentation spectrum confirming phosphorylation of NAMPT at Y188. (**K)** Quantification of NPM–ALK autophosphorylation sites identified by phosphoproteomic analysis. Phosphorylation of ALK activation sites (Y1078, Y1096, Y1131, Y1239, Y1358, Y1507, and Y1584) is reduced in cells expressing monomeric SS-AA NAMPT compared with WT NAMPT.

### NPM1::ALK and other tyrosine kinases regulate NAMPT Y188 phosphorylation in ALK⁺ ALCL

To investigate NAMPT Y188 phosphorylation in ALK⁺ anaplastic large-cell lymphoma (ALCL), we performed immunoblotting using a phospho-specific NAMPT Y188 antibody. Phosphorylation of NAMPT at Y188 was detected in most ALK⁺ ALCL cell lines (Fig. 3A) but not in ALK-ALCL cell lines. Immunohistochemical analysis of primary ALK⁺ ALCL biopsies confirmed the presence of NAMPT Y188 phosphorylation in vivo (Fig. 3B). The metabolic rewiring is a hallmark of acquired tyrosine kinase inhibitor (TKI) resistance, so we next examined NAMPT Y188 phosphorylation in ALK inhibitor (ALKi)-sensitive and resistant ALCL models. Immunoblot analysis of crizotinib and lorlatinib-resistant SUPM2 and Karpas299 derived cell lines demonstrated increased NAMPT Y188 phosphorylation relative to their parental cell lines, suggesting enhanced NAMPT activation in resistant cells (Fig. 3C). Our previous work established that the NPM–ALK–STAT3 axis transcriptionally induces NAMPT expression (*40*). We therefore hypothesized that ALKi-resistant cells further remodel NAD⁺ metabolism through coordinated regulation of NAMPT and associated substrate transporter pathways (Fig. 3D). Consistent with this model, RT–qPCR analysis showed increased NAMPT expression in lorlatinib-resistant SUPM2 and Karpas299 cells (Fig. 3E and F). Resistant cells also exhibited increased expression of the putative NMN transporter SLC12A8 (Fig. 3G and H), the mitochondrial NAD⁺ transporter SLC25A51 increased in Karpas299 cells (Fig. 3I), and in the SUPM2 cells, expression significantly decreased in lorlatinib-resistant cells (Fig. 3J).

**Fig. 3.**
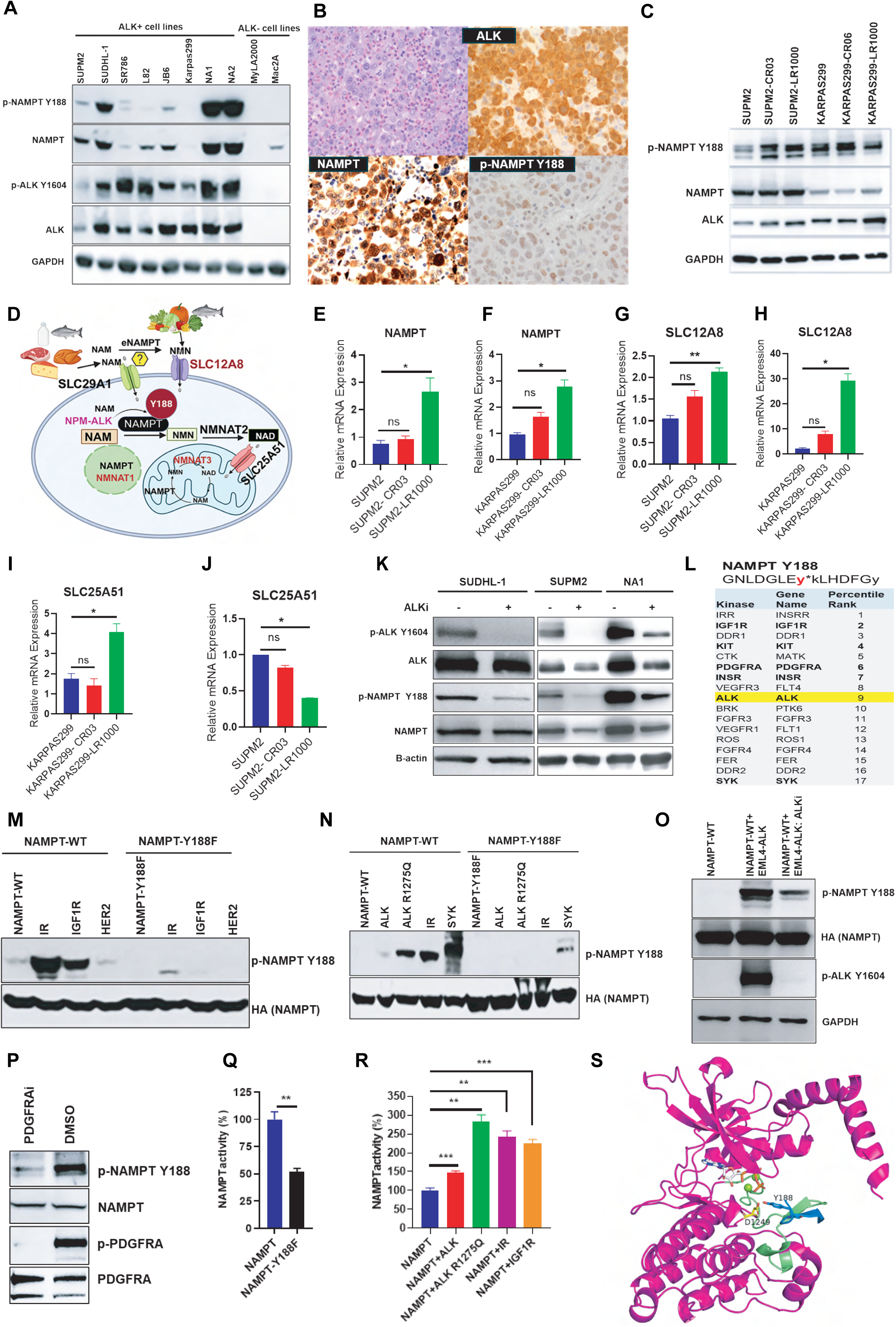
NPM1::ALK and multiple receptor tyrosine kinases regulate NAMPT Y188 phosphorylation and enzymatic activity. **(A)** Immunoblot analysis of phospho-NAMPT (Y188) and total NAMPT in ALK⁺ ALCL and mantle cell lymphoma (MCL) cell lines. (**B)** Immunohistochemical detection of phospho-NAMPT (Y188) in primary ALK⁺ ALCL patient biopsies. (**C)** Immunoblot analysis of phospho-NAMPT (Y188) in parental and ALK inhibitor-resistant SUPM2 and Karpas299 cell lines. (**D**) Schematic depicting compartmentalized NAD⁺ metabolism and transport pathways in ALKi+ALCL cells, including NAMPT, SLC12A8, SLC25A51, and SLC29A1. (**E to J)** RT–qPCR analysis of NAMPT (E and F), SLC12A8 (G and H) and SLC25A51 (I and J), (**K)** Immunoblot analysis of phospho-NAMPT (Y188) and phospho-ALK following ceritinib treatment of ALK⁺ ALCL cell lines. (**L)** Kinase prediction analysis identifying candidate kinases capable of phosphorylating NAMPT Y188. **(M)** HA immunoprecipitation and phospho-Y188 immunoblot analysis of HEK293T cells expressing HA-NAMPT-WT or HA-NAMPT-Y188F together with IR, IGF1R, or HER2. (**N)** Phospho-Y188 immunoblot analysis of HA-NAMPT-WT or HA-NAMPT-Y188F co-expressed with wild-type ALK, ALK-R1275Q, IR, or SYK. (**O)** Analysis of NAMPT Y188 phosphorylation in EML4–ALK-expressing cells treated with DMSO or lorlatinib. (**P)** Immunoblot analysis of phospho-NAMPT (Y188) in PDGFRA-transformed GIST cells treated with DMSO or avapritinib. (**Q)** Enzymatic activity of immunopurified HA-NAMPT-WT and HA-NAMPT-Y188F proteins. (**R)** NAMPT activity in cells expressing NAMPT alone or together with wild-type ALK, ALK-R1275Q, IR, or IGF1R. (**S)** Molecular docking model of a NAMPT peptide encompassing Y188 bound to the ALK kinase domain, highlighting predicted interactions with residues V1130, K1150, L1195, and L1256. Data in **E to J** are presented as mean ± s.d. from two biologically independent experiments. Statistical significance was determined using a two-tailed unpaired Student’s *t*-test as indicated in the methods.

To determine whether ALK activity directly regulates NAMPT Y188 phosphorylation, ALK⁺ ALCL cell lines (SU-DHL-1, SUPM2, and NA1) were treated with the ALK inhibitor ceritinib. Pharmacological ALK inhibition markedly reduced both ALK activation and NAMPT Y188 phosphorylation, indicating that ALK signaling contributes to this modification (Fig. 3K). To identify broader TKs capable of phosphorylating NAMPT at Y188, we performed kinase prediction analysis using https://Phosphosite.org. Among the highest-ranked candidates were ALK, insulin receptor (IR), insulin-like growth factor 1 receptor (IGF1R), insulin receptor-related receptor (INSRR), KIT, and PDGFR family members (Fig. 3L). Given the close evolutionary relationship between ALK and the insulin receptor family, we reasoned that IR and IGF1R might phosphorylate NAMPT. HEK293T cells stably expressing HA-tagged NAMPT-WT or NAMPT-Y188F were transfected with IR, IGF1R, or HER2. Immunoprecipitation followed by phospho-Y188 immunoblotting demonstrated robust phosphorylation of NAMPT-WT by IR and IGF1R, whereas HER2 showed substantially weaker phosphorylation under identical conditions (Fig. 3M). As expected, phosphorylation was abolished in the Y188F mutant, confirming site specificity. To assess the impact of oncogenic ALK signaling, we expressed wild-type ALK or the neuroblastoma-associated ALK-R1275Q mutant in NAMPT-expressing HEK293T cells. ALK-R1275Q induced markedly higher NAMPT Y188 phosphorylation than wild-type ALK, whereas phosphorylation was absent in cells expressing NAMPT alone (Fig. 3N). In parallel, SYK expression also promoted NAMPT Y188 phosphorylation, suggesting that multiple oncogenic tyrosine kinases converge on this site.

Because ALK fusion kinases are major oncogenic drivers across multiple malignancies, the EML4-ALK fusion model in Non-Small Cell Lung Cancer (NSCLC) affects the highest number of patients globally (*41*). We investigated this in our study and found that EML4–ALK increased NAMPT Y188 phosphorylation, an effect suppressed by lorlatinib treatment, confirming ALK kinase-dependent regulation of this modification (Fig. 3O). Extending these findings beyond ALK-driven cancers, PDGFRA-transformed gastrointestinal stromal tumor (GIST) cells (*42*) exhibited elevated NAMPT Y188 phosphorylation that was reduced following treatment with the PDGFRA inhibitor avapritinib, indicating that receptor tyrosine kinase-dependent regulation of NAMPT is not restricted to ALK (Fig. 3P). To determine the functional importance of Y188, HA-tagged NAMPT-WT and NAMPT-Y188F were immunopurified from HEK293T cells and subjected to enzymatic activity assays. Mutation of Y188 significantly impaired NAMPT catalytic activity, demonstrating that this residue contributes to optimal enzyme function (Fig. 3Q). Furthermore, co-expression of ALK-R1275Q, IR, or IGF1R significantly enhanced NAMPT activity compared with NAMPT alone or NAMPT co-expressed with wild-type ALK, linking Y188 phosphorylation to increased enzymatic activity (Fig. 3R). Finally, to explore the structural basis of this regulation, we performed molecular docking using a seven-residue NAMPT peptide encompassing Y188 (GLE**Y188**KLH) and the ALK kinase domain. The model predicted direct engagement of the Y188-containing motif within the ALK catalytic cleft, including hydrophobic interactions with residues V1130, K1150, L1195, and L1256. Notably, V1130 and K1150 are located within the ATP-binding region and are critical for ALK catalytic function, supporting a direct kinase–substrate interaction between ALK and NAMPT (Fig. 3S).

### NAMPT dimerization is required for Y188 phosphorylation and enzymatic activity

NAMPT functions as a homodimer to sustain NAD⁺ biosynthesis (*43*). To determine whether dimerization influences Y188 phosphorylation and catalytic activity, we generated HA–tagged NAMPT constructs encoding wild-type NAMPT (WT), a phosphorylation-deficient mutant (Y188F), and a dimerization-defective mutant. These constructs were stably expressed in the ALK⁺ ALCL cell lines SUPM2 and SUDHL1. Immunoblot analysis using phospho-specific NAMPT Y188 antibodies demonstrated complete loss of Y188 phosphorylation in the Y188F mutant and a greater than threefold reduction in the SS-AA mutant relative to WT (Fig. 4A and B). Consistent with these findings, NAMPT enzymatic activity was significantly impaired in both mutants, with Y188F and SS-AA exhibiting >50% and >90% reductions in activity, respectively (Fig. 4C). Similar effects on Y188 phosphorylation and enzymatic activity were observed in an independent ALK⁺ ALCL model, SU-DHL-1, confirming that the requirement for dimerization is not cell-line specific (Fig. 4D to F)). To directly assess the impact of Y188 phosphorylation on NAMPT dimerization, protein lysates from SUPM2 cells expressing WT, Y188F, or SS-AA NAMPT were analyzed under non-denaturing conditions by native PAGE. ACLY, a tetrameric enzyme, served as a positive control for native complex formation. Immunoblotting with anti-HA antibodies revealed that WT NAMPT migrated predominantly as a dimer, whereas the Y188F mutant retained dimer formation but with reduced abundance. In contrast, the SS-AA mutant failed to form detectable dimers, consistent with disruption of the dimerization interface (Fig. 4G). Notably, the phospho-Y188 signal was detected exclusively within the NAMPT WT dimer and was absent from Y188F-expressing cells (Fig. 4H). To show differences in protein abundance at the monomeric form of NAMPT, lysates were analyzed under denaturing conditions following HA immunoprecipitation. Comparable to WT, Y188F, and SS-AA proteins were observed in both input and immunoprecipitated fractions (Fig. 4I). Consistent with native gel analyses, phospho-Y188 signal was detected only in WT NAMPT (Fig. 4J). NPM1::ALK promotes NAMPT Y188 phosphorylation. We next tested the association between NAMPT and ALK. Co-immunoprecipitation experiments revealed that NAMPT-WT interacts with both total ALK and activated ALK (pALK-Y1604), supporting a model in which ALK preferentially associates with catalytically competent dimeric NAMPT to promote Y188 phosphorylation and enzymatic activation (Fig. 4K and L).

**Fig. 4.**
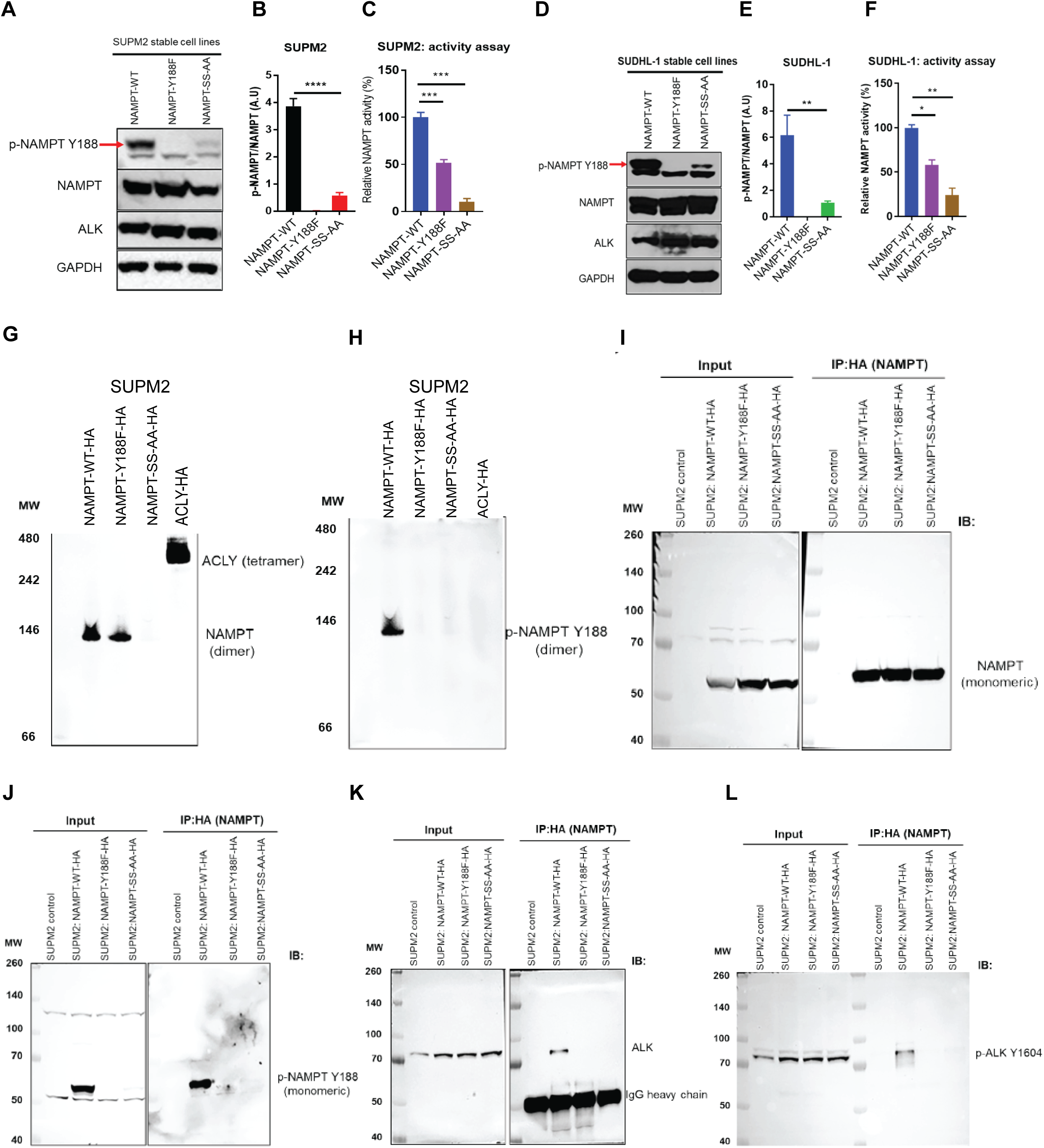
NAMPT dimerization is required for Y188 phosphorylation, enzymatic activity, and interaction with ALK. **(A)** Representative immunoblot analysis of phospho-NAMPT (Y188) and HA-tagged NAMPT in SUPM2 cells stably expressing NAMPT-WT, NAMPT-Y188F, or the dimerization-deficient NAMPT-SS-AA (S199A/S200A) mutant. (**B)** Quantification of phospho-NAMPT (Y188) normalized to total HA-NAMPT. (**C)** NAMPT enzymatic activity measured in immunopurified WT, Y188F, and SS-AA proteins from SUPM2 cells. (**D)** Representative immunoblot analysis of phospho-NAMPT (Y188) and HA-NAMPT in SU-DHL-1 cells expressing WT, Y188F, or SS-AA NAMPT. (**E)** Quantification of phospho-NAMPT (Y188) in SU-DHL-1 cells. (**F)** NAMPT enzymatic activity measured in SUDHL-1-derived samples. (**G)** Native PAGE analysis of HA-tagged NAMPT complexes from SUPM2 cells expressing WT, Y188F, or SS-AA NAMPT. ACLY served as a positive control for higher-order oligomer formation. (**H)** Native PAGE immunoblot probed with phospho-NAMPT (Y188) antibody demonstrating phosphorylation of dimeric WT NAMPT. (**I),** Denaturing SDS–PAGE analysis of input and HA-immunoprecipitated fractions showing comparable expression and recovery of WT, Y188F, and SS-AA NAMPT proteins. (**J)** Immunoblot analysis of phospho-NAMPT (Y188) in HA-immunoprecipitated fractions. (**K and L),** Co-immunoprecipitation analysis demonstrating association of WT NAMPT with total ALK (**K**) and activated ALK (pALK-Y1604) (**L**). Data in **B, C, E,** and **F** are presented as mean ± s.d. from three biologically independent experiments. Statistical significance was determined using two-tailed unpaired Student’s *t*-test, as described in the methods.

### NAMPT and NPM1::ALK co-localize in subcellular compartments, including mitochondria

While NPM1::ALK is known to localize in cytoplasm and nucleus, several oncogenic tyrosine kinases, including c-Abl, ERBB2, SRC, and FGFR1, have been reported to localize to mitochondria, where they regulate metabolic and survival pathways (*44–47*). To delineate in greater depth interactions of NPM1::ALK and NAMPT, we fractionated cellular lysates into cytosolic, mitochondrial, and nuclear compartments of SUPM2 cells, which stably express WT, Y188F mutant, or NAMPT SS-AA monomer. Comparative western blot analysis of the cytosolic and nuclear fractions using antibodies against total and phosphorylated NAMPT and NPM1::ALK revealed that NPM1::ALK and NAMPT-WT, including their active, phosphorylated forms, are present in both compartments, with the majority residing in the cytosol (Fig. 5A and B). The inability of the NAMPT-SS-AA mutant to form dimers not only affected its phosphorylation at Y188 but also its expression and localization in both these compartments. Comparative analysis of the mitochondrial fraction (Fig. 5C and D) also detected the presence of NPM1::ALK and NAMPT-WT, total and phosphorylated for both, in concentrations akin to that seen in the nucleus. Mitochondrial expression of the NAMPT-SS-AA mutant was impaired, similarly to its nuclear localization. Notably, introduction of the NAMPT-Y188F mutant markedly augmented the expression and phosphorylation of endogenous NAMPT and, strikingly, NPM1::ALK. This finding suggests an active functional interaction between NPM1::ALK and NAMPT, with the increase in expression of the latter induced by the former, most likely aimed at maintaining adequate NAD+ supply within mitochondria, adversely affected by co-expression of the hypofunctional NAMPT-Y188F mutant. A schematic model summarizing NAMPT localization and trafficking across cellular compartments is shown (Fig. 5E). To determine whether these observations extend to primary disease samples, we performed immunofluorescence and confocal microscopy on ALK⁺ ALCL patient specimens using antibodies against ALK, NAMPT, and the mitochondrial marker VDAC1. Triple-labelling analysis revealed co-localization of both ALK and NAMPT with mitochondrial structures marked by VDAC1 (Fig. 5F). Higher-magnification images further demonstrated the spatial overlap of ALK and NAMPT within mitochondria, supporting the existence of a mitochondrial ALK–NAMPT signaling axis in ALK-driven lymphoma (Fig. 5F, right). Together, these findings identify mitochondria as a previously unrecognized subcellular site of both NAMPT and NPM1::ALK localization in ALK⁺ ALCL and suggest that NAMPT may facilitate compartment-specific trafficking that contributes to localized NAD⁺ metabolism. Notably, mitochondrial localization of NAMPT has previously been reported in HEK293T cells, supporting our observations (*48*). In contrast, another study in HeLa cells did not detect mitochondrial NAMPT localization (*49*). These discrepant findings suggest that mitochondrial localization of NAMPT may be context-dependent, varying according to cell type, oncogenic state, or metabolic adaptations that support the elevated NAD⁺ requirements of tumor cells.

**Fig. 5.**
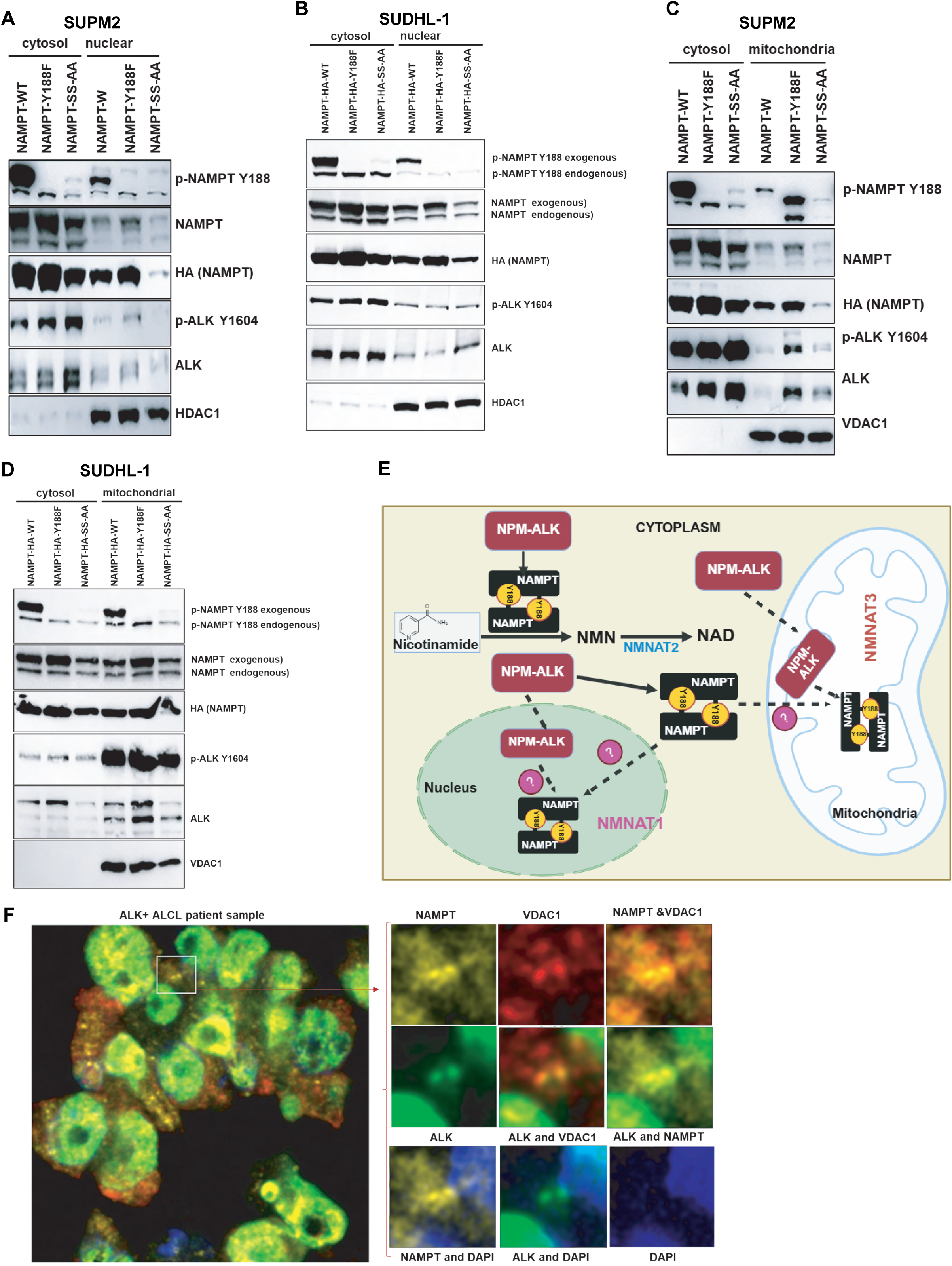
NAMPT dimerization promotes mitochondrial and nuclear localization, and NPM1::ALK co-localizes with NAMPT in mitochondria. **(A and B)** Subcellular fractionation and immunoblot analysis of cytosolic and mitochondrial fractions from SUPM2 and SUDHL-1 cells stably expressing HA-tagged NAMPT-WT, NAMPT-Y188F, or the dimerization-deficient NAMPT-SS-AA (S199A/S200A) mutant. Immunoblots were probed with the indicated antibodies to assess NAMPT localization. Fraction-specific markers were used to verify sample purity. **(C and D)** Immunoblot analysis of cytosolic and nuclear fractions from SUPM2 and SUDHL-1 cells expressing NAMPT-WT, NAMPT-Y188F, or NAMPT-SS-AA, demonstrating nuclear accumulation of dimeric NAMPT. Fraction purity was confirmed using compartment-specific marker proteins. **(E)** Schematic model illustrating NAMPT trafficking between cytosolic, mitochondrial, and nuclear compartments and the proposed role of dimerization in regulating subcellular localization and compartmentalized NAD⁺ metabolism. **(F),** Representative confocal immunofluorescence images of ALK⁺ ALCL patient samples stained for NAMPT, ALK, and the mitochondrial marker VDAC1. Right, higher-magnification view of the boxed region highlighting mitochondrial co-localization.

### NPM1::ALK triggers NAMPT-dependent NAD+ biosynthesis

Having established that Y188 phosphorylation enhances NAMPT enzymatic activity, we next examined whether ALK signaling regulates NMN and NAD metabolism. To address this question, we performed targeted metabolomic profiling using liquid chromatography–tandem mass spectrometry (LC–MS/MS) in ALK inhibitor-sensitive (ALKi-S) and lorlatinib-resistant (ALKi-R) SUPM2 ALK⁺ ALCL cells following treatment with either DMSO or lorlatinib as shown schematically in Fig. 6A and Fig.6B, respectively. To validate metabolite identification, the LC–MS/MS platform was calibrated using authentic standards for NAD⁺, NADH, NADP⁺, NADPH, and NMN (Fig. S1A and B). Metabolomic analysis revealed widespread alterations in metabolites associated with the NAMPT-dependent NAD⁺ biosynthetic pathway following ALK inhibition. ATP accumulated in ALKi-S cells following lorlatinib treatment, it remained unchanged in ALKi-R cells (Fig. 6C and D), consistent with inhibition of NPM1::ALK activity in the former. Because nicotinamide mononucleotide (NMN) is the direct product of NAMPT, we next examined NMN abundance. While lorlatinib treatment profoundly reduced NMN level in ALKi-S, it was much less reduced in ALKi-R cells (Fig. 6E and F). Accordingly, ALK inhibition resulted in a profound reduction in NAD⁺ levels in ALKi-S (Fig. 6G), the NAD⁺ abundance fully maintained in ALKi-R cells, suggesting possible NMN-independent supplementation of NAD+ synthesis (Fig. 6H). These findings support the existence of adaptive metabolic mechanisms that sustain NAD⁺ metabolism in the context of acquired ALK inhibitor resistance from nicotinic acid (Preiss–Handler pathway) and QPRT/tryptophan (de novo/kynurenine pathway) (*24, 50*). Consistent with the NAD+ reduction in ALKi-S cells, AMP levels were significantly decreased in ALKi-S cells following lorlatinib treatment, whereas no significant changes were observed in the resistant cells (Fig. 6I and J). To validate these metabolomic findings, intracellular NAD⁺ levels were quantified after 48 h treatment with the ALKi lorlatinib and the NAMPTi FK866, either alone or in combination, in ALKi-S and ALKi-R SUP-M2 and Karpas299 cells using a commercially available NAD⁺ assay kit. Consistent with the metabolomic analysis, lorlatinib treatment significantly reduced NAD⁺ levels in ALKi-S cells but had little effect on NAD⁺ abundance in ALKi-R cells. In contrast, FK866 treatment markedly depleted intracellular NAD⁺ levels in both sensitive and resistant cells, confirming effective inhibition of NAMPT activity. Furthermore, combined treatment with lorlatinib and FK866 produced an even greater reduction in NAD⁺ levels in both cell lines compared with either treatment alone, demonstrating that pharmacological inhibition of NAMPT effectively overcomes the maintenance of NAD⁺ pools observed in ALKi-resistant cells (Fig.6K and L). Together, these data demonstrate that NPM1::ALK signaling regulates NAMPT-dependent NMN production and contributes to the maintenance of cellular NAD⁺ pools.

**Fig. 6.**
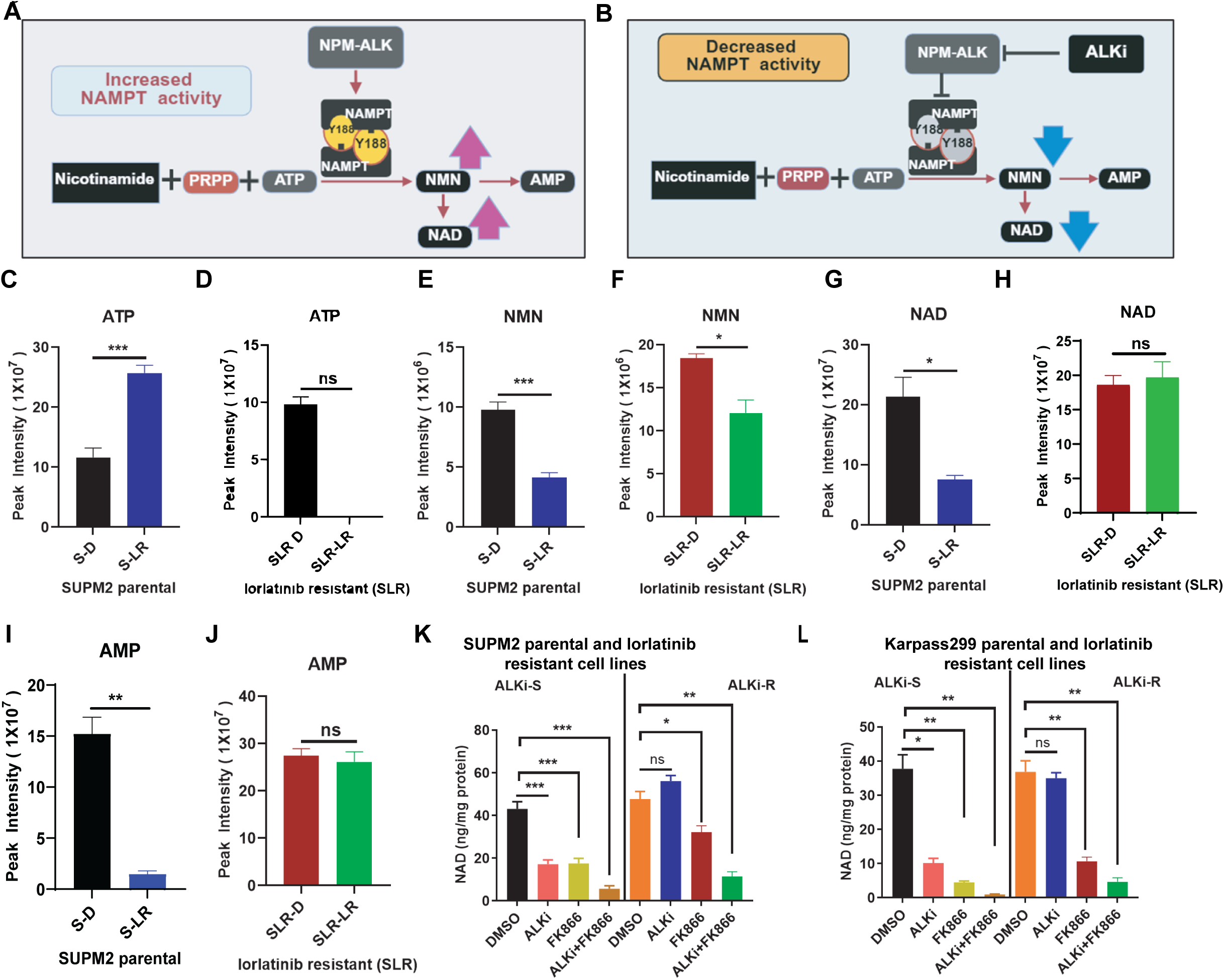
NPM1::ALK signaling regulates NAMPT-dependent NMN and NAD metabolism. **(A)** Schematics of targeted LC–MS/MS metabolomic profiling of ALK inhibitor-sensitive (ALKi-S) SUPM2 cells treated with DMSO or 50 nM lorlatinib for 24 h. (**B)** Experimental workflow for metabolomic profiling of lorlatinib-resistant (ALKi-R) SUPM2 cells treated with DMSO or 200 nM lorlatinib for 24 h. **(C and D)** Intracellular ATP levels in ALKi-S (**C**) and ALKi-R (**D**) cells. **(E and F)** Quantification of nicotinamide mononucleotide (NMN), the direct product of NAMPT, in ALKi-S (**E**) and ALKi-R (**F**) cells. **(G and H)** Intracellular NAD⁺ levels in ALKi-S (**G**) and ALKi-R (**H**) cells. **(I and J)** AMP abundance in ALKi-S (**I**) and ALKi-R (**J**) cells. (**K)** Parental SUPM2, ALKi-S and lorlatinib-resistant SUPM2-LR1000 (ALKi-R) cells were treated with DMSO, lorlatinib (ALKi-S, 50 nM and ALKi-R, 200 nM), FK866 (25 nM), alone or the combination for 48 h, and NAD+ was measured by using commercially available kit. **(L),** Parental Karpas299, ALKi-S and lorlatinib-resistant Karpas299-LR1000 (ALKi-R) cells were treated with DMSO, lorlatinib (ALKi-S, 50 nM and ALKi-R, 300 nM), FK866 (25 nM), alone or the combination for 48 h, and NAD+ was measured by using commercially available kit. Targeted metabolite measurements were performed by LC–MS/MS following 24 h treatment with DMSO or lorlatinib. Data are presented as mean ± s.d. from three biologically independent experiments. Statistical significance was determined using two-tailed unpaired Student’s *t*-test. The PRISM GraphPad software was used to analyze the t-test (p> value; 0.05 *, 0.005 ** 0.0005 ***).

### NAMPT monomer/dimer and phosphorylation remodel distinct protein-protein interaction networks

To define how NAMPT oligomerization (monomer/dimer) and tyrosine phosphorylation influence its interactome, we performed LC–MS/MS analysis on immunoprecipitation (IP) of HA-tagged dimeric NAMPT-WT and monomeric NAMPT-SS-AA in the presence or absence of oncogenic NPM1::ALK signaling. Following normalization to vector controls and stringent filtering, 8,303 high-confidence proteins were identified across all conditions. Principal component analysis revealed clear separation among the NAMPT-WT, NAMPT-WT+ALK, NAMPT-SS-AA, and NAMPT-SS-AA+ALK interactomes, indicating that both oligomerization state and ALK-dependent phosphorylation substantially reshape NAMPT-associated protein networks (Fig. 7A). Comparable protein abundance distributions and protein identification rates across samples confirmed the quality and reproducibility of the dataset (Fig. 7B and C). Consistent with extensive interactome remodeling, unsupervised clustering identified six protein modules displaying distinct abundance patterns across the four conditions (Fig. 7D). Using a threshold of log2 fold change ≥1 relative to vector controls, we identified 1,798, 2,207, 1,963 and 1,645 enriched proteins in the NAMPT-WT, NAMPT-WT+ALK, NAMPT-SS-AA and NAMPT-SS-AA+ALK interactomes, respectively (Fig. 7E). Comparative analysis revealed both shared and condition-specific interaction networks, suggesting that oligomerization and phosphorylation independently contribute to NAMPT interactome organization. Pathway enrichment analysis of proteins uniquely associated with dimeric NAMPT-WT identified metabolic processes, including carbon metabolism, fatty acid metabolism, propanoate metabolism and the tricarboxylic acid cycle, consistent with the established metabolic functions of NAMPT. Phosphorylation of dimeric NAMPT by NPM1::ALK markedly expanded this network, resulting in enrichment of pathways linked to amino acid biosynthesis and endocytosis in addition to core metabolic programs (Fig. 7F). In contrast, proteins uniquely associated with monomeric NAMPT-SS-AA were enriched for pathways involved in base excision repair and cell-cycle regulation, supporting a functional shift toward nuclear processes. Notably, ALK-dependent phosphorylation redirected the monomeric NAMPT interactome toward pathways associated with ribosome biogenesis, ribosomal function and RNA transport, indicating extensive rewiring of RNA-processing and translational machinery interactions (Fig. 7G). Despite these state-dependent changes, 912 proteins were shared across all four conditions. This conserved interactome was enriched for oxidative phosphorylation, metabolic pathways and kinase-associated signaling networks, suggesting the presence of a core NAMPT interaction scaffold that is maintained independently of oligomerization status or phosphorylation state.

**Fig. 7.**
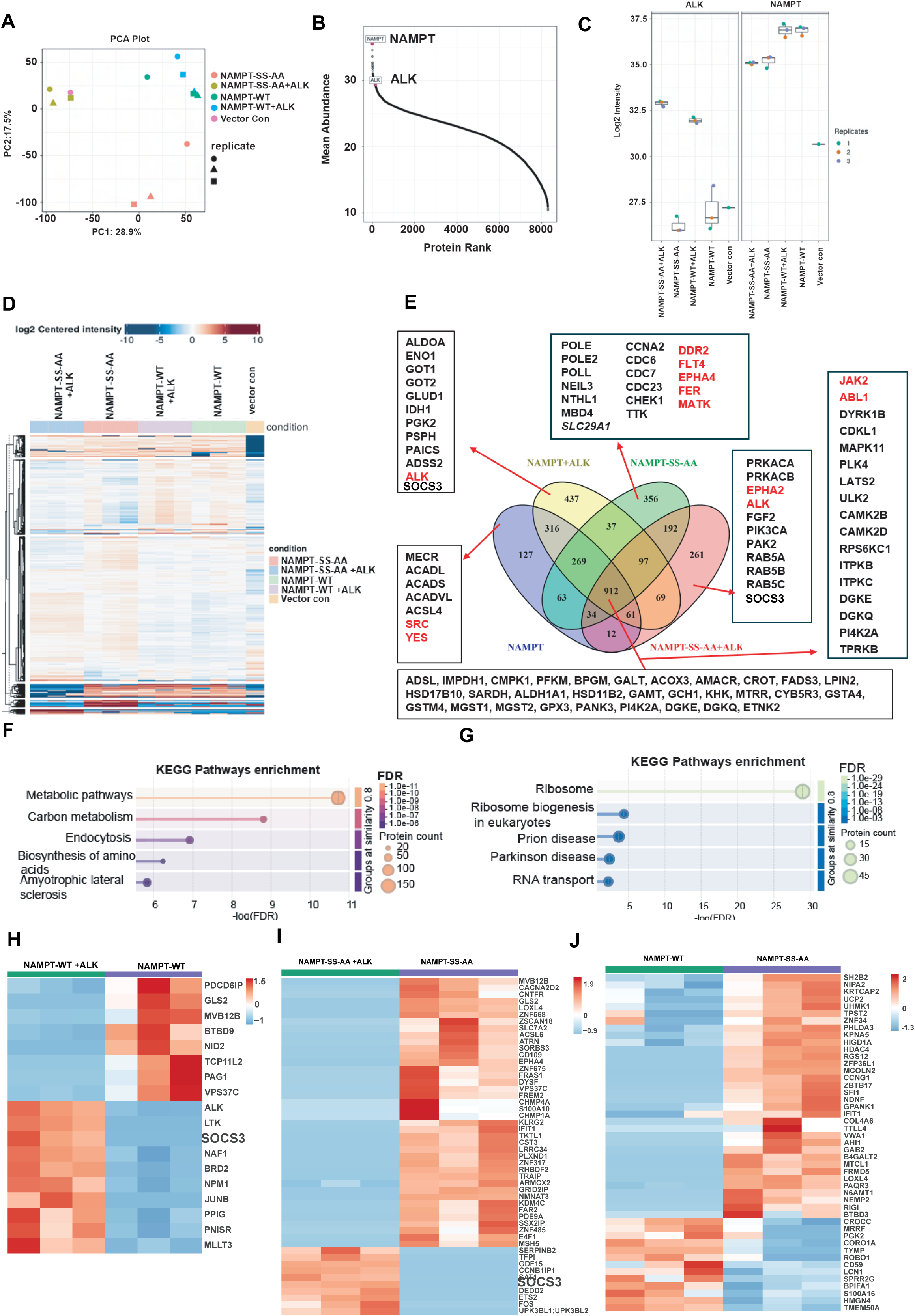
NAMPT oligomerization and tyrosine phosphorylation remodel distinct protein–protein interaction networks. **(A)** Principal component analysis (PCA) of LC–MS/MS datasets obtained from HA-tag immunoprecipitation of NAMPT-WT and NAMPT-SS-AA in the presence or absence of NPM–ALK, showing distinct clustering of the four interactomes. **(B and C)** Quality-control analyses showing protein abundance distributions and numbers of proteins identified across samples. (**D)** Unsupervised hierarchical clustering of enriched interactors identifies six protein modules with distinct abundance patterns across experimental conditions. **(E),** Venn diagram showing overlap and condition-specific proteins enriched relative to vector controls (log2 fold change ≥ 1). A total of 1,798, 2,207, 1,963, and 1,645 proteins were enriched in the NAMPT-WT, NAMPT-WT+ALK, NAMPT-SS-AA, and NAMPT-SS-AA+ALK interactomes, respectively, with 912 proteins shared across all conditions. **(F)** KEGG pathway enrichment analysis of proteins uniquely associated with NAMPT-WT and NAMPT-WT+ALK interactomes. (**G)** KEGG pathway enrichment analysis of proteins uniquely associated with NAMPT-SS-AA and NAMPT-SS-AA+ALK interactomes. (**H)** Differential interactome analysis comparing NAMPT-WT+ALK and NAMPT-WT identifies proteins preferentially associated with phosphorylated or non-phosphorylated dimeric NAMPT (absolute fold change ≥ 2; adjusted P < 0.05). (**I)** Differential interactome analysis of NAMPT-SS-AA+ALK versus NAMPT-SS-AA showing phosphorylation-dependent changes in the monomeric NAMPT interactome. (**J)** Direct comparison of dimeric and monomeric NAMPT interactomes reveals extensive differences in protein association patterns and a broader interaction landscape for monomeric NAMPT. Differentially enriched proteins in h–j were defined using an absolute fold change ≥ 2 and adjusted *P* < 0.05. KEGG pathway enrichment analyses were performed using proteins uniquely detected in each condition.

Comparison of the NAMPT-WT+ALK and NAMPT-WT interactomes identified a discrete set of proteins selectively enriched with phosphorylated dimeric NAMPT (absolute fold change ≥2; adjusted *P* < 0.05), including ALK, LTK, SOCS3, BRD2 and JUNB, whereas PAG1, GLS2, NID2 and BTBD9 preferentially associated with non-phosphorylated NAMPT, indicating that tyrosine phosphorylation remodels the dimeric NAMPT interactome (Fig. 7H). In contrast, monomeric NAMPT-SS-AA displayed a broader interactome, with substantially more proteins associated with the non-phosphorylated than the phosphorylated form, suggesting that tyrosine phosphorylation restricts interaction networks accessible to monomeric NAMPT (Fig. 7I). SOCS3 was enriched in both phosphorylated dimeric and monomeric NAMPT interactomes, identifying it as a shared phospho-dependent NAMPT interactor. Direct comparison of dimeric and monomeric NAMPT further revealed extensive differences in binding partners, with the monomeric species exhibiting a broader interaction landscape (Fig. 7J).

Comparative interactome profiling identified distinct protein interaction landscapes associated with ALK expression and NAMPT phosphorylation status. Differentially enriched interactors in the NAMPT-WT versus NAMPT-WT+ALK, NAMPT-SS-AA versus NAMPT-SS-AA+ALK, NAMPT-WT+ALK versus NAMPT-SS-AA+ALK, and NAMPT-WT versus NAMPT-SS-AA comparisons are shown in volcano plots (Fig. S2A to D).

### Tyrosine phosphorylation of NAMPT by NPM1::ALK promotes recruitment of SOCS3

Although dimeric NAMPT was substantially more abundant than monomeric NAMPT in both the immunoprecipitated bait protein and proteomic datasets (Fig. S3A), NPM1::ALK preferentially associated with Y188-phosphorylated monomeric NAMPT, indicating that this interaction is selective and independent of protein abundance (Fig. S3B). Since Y188 resides at the NAMPT dimer interface, disruption of dimerization by the SS-AA mutant is predicted to increase Y188 accessibility, facilitating phosphotyrosine-dependent binding by NPM1::ALK. Similarly, SOCS3 is an SH2 domain-containing adaptor that recognizes phosphotyrosine-containing proteins (*51*), preferentially associated with Y188-phosphorylated monomeric NAMPT despite its low abundance (Fig. S3C). In addition, the epigenetic reader BRD2 was strongly enriched in the p-NAMPT Y188 interactome compared with non-phosphorylated NAMPT (Fig. S3D). STRING analysis of proteins enriched in the p-NAMPT Y188 interactome (log₂ fold change >2, *P* < 0.05) identified a prominent ALK–NAMPT–SOCS3 signaling network (Fig. S3E and F), supporting the biochemical interactions observed by proteomics. Public datasets from the Human Protein Atlas further revealed high SOCS3 expression in ALK⁺ ALCL cell lines (Fig. S3G). Comparison with the BioGRID database identified 28 shared NAMPT interactors, including MYC, KRAS, BRD4 and BRD2, while uncovering more than 1,484 previously unreported NAMPT-associated proteins (Fig. S3H). Co-immunoprecipitation confirmed BRD4 association with both phosphorylated and non-phosphorylated NAMPT (Fig. S3I). Integrated STRING analysis of p-NAMPT Y188, monomeric and dimeric interactomes showed that Y188 phosphorylation preferentially enriches phosphoprotein-associated signaling networks, whereas monomeric and dimeric NAMPT engage distinct metabolic and regulatory modules (Fig. S4 A to D), positioning NAMPT as a central node linking NAD⁺ metabolism to signaling and stress adaptation.

### NAMPT Y188 phosphorylation and dimerization are critical for cell growth

To assess the functional consequences of NAMPT Y188 phosphorylation on cell proliferation, we cultured SUPM2 cells stably expressing NAMPT-HA-GFP-WT, NAMPT-HA-GFP-Y188F, and NAMPT-HA-GFP-SS-AA in 96-well plates (5 × 10^3^ cells/well) for 72 hours. Cell growth was monitored via green fluorescence intensity (N=6). The results showed that cells expressing the phosphorylation-deficient NAMPT-Y188F mutant exhibited over a 45% reduction in proliferation (p < 0.001) compared to NAMPT-WT-expressing cells, while those expressing the dimerization-deficient NAMPT-SS-AA mutant showed more than a 35% decrease (p < 0.01) (Fig. 8A). When we evaluated the clonogenic potential of SUPM2 cells via methylcellulose colony formation assays, cells expressing NAMPT-WT formed significantly more colonies by ∼70% (p < 0.005) relative to cells expressing either NAMPT-Y188F or NAMPT-SS-AA mutant (Fig. 8B and C). Next, we evaluated the effects of the ALKi, lorlatinib and the NAMPTi FK866, applied either alone or in combination, on the proliferation of ALKi-S and ALKi-R SUPM2 and Karpas299 cells following 72 h of treatment. As expected, lorlatinib significantly reduced cell proliferation in ALKi-S cells but had no significant effect on ALKi-R cells. In contrast, FK866 significantly attenuated cell proliferation in both ALKi-S and ALKi-R cell lines. Moreover, combined treatment with lorlatinib and FK866 resulted in a greater reduction in cell proliferation than either agent alone in both cell lines (Fig. 8D and E). These findings demonstrate that pharmacological inhibition of NAMPT effectively overcomes ALK inhibitor resistance and suppresses the growth of ALKi-resistant lymphoma cells. Together, these findings support a model in which NPM1::ALK phosphorylates NAMPT at Y188, thereby enhancing NMN and NAD+ biosynthesis and promoting cancer cell growth (Fig.8F).

**Fig. 8.**
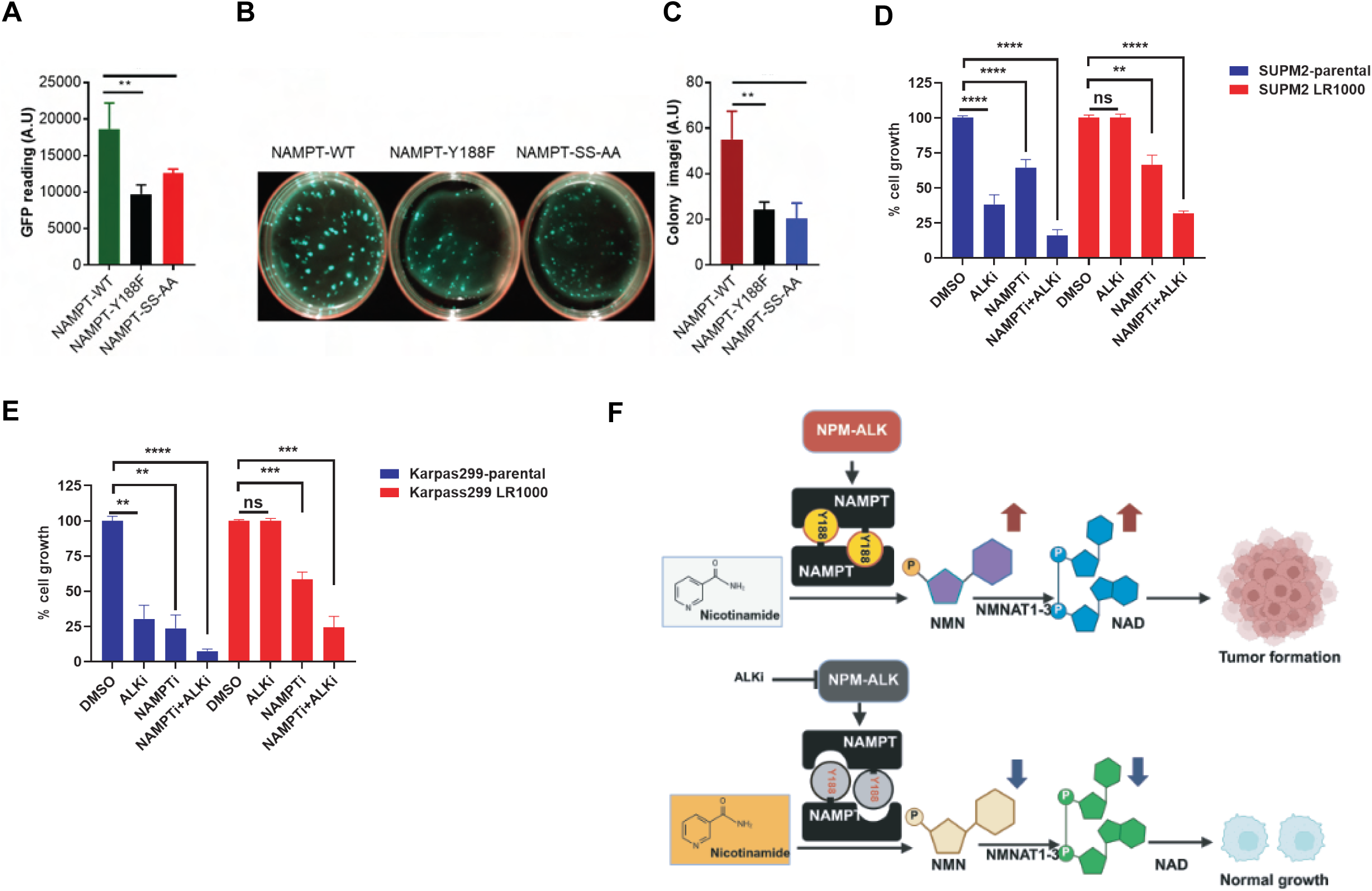
|NAMPT Y188 phosphorylation promotes NAMPT-dependent proliferation and sensitizes lorlatinib-resistant ALCL cells to combined NAMPT and ALK inhibition. **(A)** Proliferation of SUPM2 cells expressing NAMPT-WT, the phosphorylation-deficient mutant Y188F, or the dimerization-deficient mutant SS-AA was quantified by GFP fluorescence over 72 h. Both Y188F and SS-AA significantly reduced proliferation relative to WT. **(B and C)** Representative images (B) and quantification **(C)** of methylcellulose colony formation assays showing reduced clonogenicity of Y188F- and SS-AA-expressing cells compared with WT. (**D)** Parental SUPM2, ALKi-S and lorlatinib-resistant SUPM2-LR1000 (ALKi-R) cells were treated with DMSO, lorlatinib (ALKi-S, 50 nM and ALKi-R, 200 nM), FK866 (25 nM), alone or the combination for 72 h, and proliferation was measured by WST-1 assay. (**E)** Parental Karpas299, ALKi-S and lorlatinib-resistant Karpas299-LR1000 (ALKi-R) cells were treated with DMSO, lorlatinib (ALKi-S, 50 nM and ALKi-R, 300 nM), FK866 (25 nM), alone or the combination for 72 h, and proliferation was measured by WST-1 assay. (**F)** A schematic illustrating that oncogenic tyrosine kinase-mediated phosphorylation of NAMPT at Y188 enhances NAMPT activity, NAD⁺ biosynthesis, and cancer cell proliferation. Data are presented as mean ± s.d. from three biologically independent experiments. Statistical significance was determined using two-tailed unpaired Student’s *t*-test. The PRISM GraphPad software was used to analyze the t-test (p> value; 0.05 *, 0.005 **, 0.0005 ***, 0.0001****).

## DISCUSSION

NAMPT is the rate-limiting enzyme in the NAD+ salvage pathway, placing it at the center of cellular metabolism, stress responses, and signaling in both physiological and pathological contexts, including cancer (*52, 53*). Although the mechanisms regulating NAMPT expression have been extensively investigated (*54*), those that directly control its enzymatic activity remain poorly defined. Here, we identify tyrosine phosphorylation as a previously unrecognized post-translational mechanism that activates NAMPT, revealing a new mode of regulation of NAD+ biosynthesis. We show that multiple oncogenic tyrosine kinases, including NPM1::ALK, EML4::ALK, IGF1R, INSR and PDGFRA, phosphorylate NAMPT, with phosphorylation of Y188 enhancing enzymatic activity, NAD+ biosynthesis, downstream metabolic outputs and cell growth. These findings establish a direct mechanistic link between oncogenic kinase signaling and NAD+ synthesis, suggesting that oncogenic kinases sustain the metabolic demands of malignant cells by activating the rate-limiting enzyme of the NAD+ salvage pathway. Consistent with this model, the neuroblastoma-associated ALK-R1275Q mutant promoted NAMPT phosphorylation and catalytic activation more efficiently than wild-type ALK, suggesting that enhanced stimulation of NAD⁺ biosynthesis is likely a conserved feature of oncogenic ALK variants (*55, 56*). More broadly, identifying NAMPT as a substrate of several oncogenic tyrosine kinases suggests that phosphorylation-dependent regulation of NAD⁺ metabolism may represent a common mechanism supporting tumor growth across genetically diverse malignancies.

Our data further demonstrates that NAMPT dimerization is required for efficient Y188 phosphorylation and maximal enzymatic activity, providing a structural basis for kinase-dependent regulation. Consistent with structural studies showing that the active site is assembled at the dimer interface, where Y188 lies adjacent to catalytic residues (*57*), disruption of dimerization by the SS-AA substitution markedly reduced Y188 phosphorylation and catalytic activity, suggesting that phosphorylation may stabilize an active conformation or modulate local structural dynamics. These observations indicate that kinase-mediated regulation of NAMPT is tightly coupled to its structural organization, providing an additional level of control over NAD⁺ biosynthesis.

Beyond catalysis, oligomerization and phosphorylation extensively remodeled the NAMPT interactome. Whereas dimeric NAMPT preferentially associated with metabolic proteins, monomeric NAMPT interacted with proteins involved in DNA repair, cell-cycle regulation and RNA processing. Phosphorylation further rewired these interaction networks, revealing previously unrecognized signaling connections. Notably, SOCS3 associated with phosphorylated NAMPT irrespective of its oligomeric state, indicating that phosphotyrosine-dependent interactions can occur independently of structural organization. Together, these findings identify NAMPT as both a metabolic enzyme and a dynamic signaling node whose functions are coordinated by dimerization and phosphorylation. Phosphorylated NAMPT dimer (p-NAMPT-WT) and phosphorylated NAMPT-SS-AA monomer defined two distinct Hallmark-enriched cellular programs with clear functional separation in both signaling and metabolism. The enrichment of antioxidant enzymes, including PRDX1/2/4, SOD1/2, TXNRD2, and GSR, together with TCA cycle-associated enzymes such as GOT1/2, GLUD1, and IDH1, suggests enhanced redox buffering and mitochondrial metabolic plasticity, features increasingly recognized as essential for metastatic progression and adaptation to oxidative stress (*58–60*). In contrast, phosphorylated NAMPT-SS-AA monomer was dominated by MYC target pathways, ribosome biogenesis, nucleolar organization, and translational machinery. Extensive enrichment of ribosomal proteins, aminoacyl-tRNA synthetases, translation initiation factors, ER-Golgi trafficking proteins, heat-shock proteins, and proteasome components indicates a highly anabolic cellular program characterized by elevated protein synthesis and proteostasis demand, hallmarks of MYC-driven tumors (*61, 62*). Concurrent enrichment of glycolytic enzymes, including PKM, LDHA, and SLC2A1, further supports metabolic reprogramming toward aerobic glycolysis to sustain rapid biomass accumulation (*63, 64*). Together, these observations indicate that phosphorylation and oligomerization cooperate to define fundamentally distinct biological outputs, distinguishing an oxidative, mesenchymal-like metabolic state from a MYC-driven anabolic proliferative state. NAMPT interactome represents a high NAD⁺-turnover state in which continuous NAMPT-mediated NAD⁺ salvage is essential to sustain oxidative phosphorylation, lipid metabolism, and genome maintenance (*65, 66*).

Our findings also have therapeutic implications. Although NAMPT inhibitors have shown promise in preclinical models, their clinical development has been limited by toxicity and an incomplete understanding of tumor dependency on NAMPT-driven NAD⁺ metabolism (*67, 68*). Our results demonstrate that pharmacological inhibition of NAMPT effectively suppresses the growth of both ALK inhibitor-sensitive and ALK inhibitor-resistant lymphoma cells, and that combined treatment with lorlatinib and FK866 produces greater growth inhibition than either agent alone (Fig.8). These findings suggest that cancers driven by ALK and potentially other oncogenic tyrosine kinases remain dependent on phosphorylation-mediated NAMPT activation even after acquisition of kinase inhibitor resistance. Consequently, phospho-Y188 NAMPT may represent both a biomarker of elevated NAD⁺ salvage activity and a predictor of response to metabolic therapies targeting the NAD⁺ salvage pathway. Targeting kinase-mediated NAMPT activation, disrupting dimerization, or selectively interfering with phosphorylation-dependent NAMPT functions may therefore complement catalytic NAMPT inhibition while potentially improving therapeutic selectivity.

Collectively, our study identifies tyrosine phosphorylation as a previously unrecognized layer of NAMPT regulation and establishes a direct connection between oncogenic kinase signaling, NAMPT structural organization and NAD+ metabolism. These findings position NAMPT as a signaling-responsive metabolic enzyme that integrates oncogenic inputs to coordinate metabolic adaptation and tumor cell fitness.

## MATERIALS AND METHODS

### Reagents and antibodies

CycLex^®^ NAMPT Colorimetric Assay Kit Ver.2 from MBL International (Catalog#CY-1251V2), beta-nicotinamide mononucleotide (NMN, catalog#N3501), beta-nicotinamide adenine dinucleotide (NAD catalog#N0632) standards from Sigma for validation of NMN and NAD in our metabolomics data. All the other reagents are from Sigma-Aldrich. Cell Proliferation Reagent WST-1 (Millepore-sigma, catalog# 5015944001), HA Tag Monoclonal Antibody, Pierce IP Lysis Buffer, Pierce Anti-HA Magnetic Beads, Halt Protease and Phosphatase Inhibitor, All the active Recombinant Human tyrosine kinases such as ALK (catalog# PV3867), SRC (catalog# P3044), FYN (catalog# P3042), LCK (catalog#PV4081), BTK (catalog#PV3587), FLT3 (catalog# PV3967), ABL (catalog# P3049), SYK (catalog# PV3857) and EGFR (catalog#PV4190), non-tyrosine kinases AKT1 (catalog#PR3878D) and ERK1 (catalog#PV3594) were purchased from Thermo Fisher Scientific. Recombinant human Insulin Receptor protein from Abcam (catalog#ab80251), GAPDH (catalog# 60004-1-Ig) from Proteintech, HA-Tag (C29F4) Rabbit (catlog# 3724S), Phospho-ALK (Tyr1604) Antibody (catalog# 3341S), ALK (D5F3^®^) XP^®^ Rabbit (catalog# 3633S) from Cell Signaling Technology. NAMPT activity assay kit from MBL International (catalog#CY-1251V2), NAMPT antibody (Catalog # 66385-1-Ig) from Proteintech. Insulin Receptor Beta Polyclonal Antibody (catalog#A303-712A) from Thermofisher, IGF-I Receptor β Antibody (catalog#3027S), Phospho-IGF-I Receptor β (Tyr1131)/Insulin Receptor β (Tyr1146) Antibody (catalog#3021S) from Cell Signaling Technology. NAD/NADH Assay Kit (Colorimetric) from Abcam (catalog# ab65348).

### Phospho-NAMPT Y188 custom antibody production

To validate the phosphorylation of NAMPT at tyrosine 188 (Y188) in a cellular system, we developed a custom phospho-specific antibody by immunizing two rabbits with the synthetic phosphopeptide C-NLDGLE(pY)KLHDFG-amide, followed by affinity purification. This custom antibody development was conducted in collaboration with Thermo Fisher Scientific, Inc.

### NAMPT constructs with HA-GFP-tagged wild-type or Y-F mutant plasmids and lentivirus production

To study the role of specific tyrosine residues and dimerization in NAMPT function, we generated a series of custom plasmids encoding HA-GFP-tagged NAMPT constructs. These included the wild-type NAMPT sequence and the following tyrosine-to-phenylalanine (Y→F) mutants: Y34F, Y54F, Y103F, Y188F, Y195F, and Y453F, as well as a NAMPT-SS→AA dimerization-defective mutant. All constructs were cloned into the pLenti_MS2-P65-HSF1_GFP backbone with a C-terminal HA tag. Plasmid synthesis and sequence validation were performed by GenScript (Piscataway, NJ, USA). Lentiviral particles were generated using these constructs for stable expression in mammalian cells.

### Transfection and HA-Tagged NAMPT Immunoprecipitation

HEK293T cells were transiently transfected with the above NAMPT-HA constructs using a standard lipid-based transfection method. At 48 hours post-transfection, cells were harvested and lysed in NP-40 lysis buffer supplemented with protease and phosphatase inhibitors. The HA-tagged NAMPT proteins were immunoprecipitated from cleared lysates using anti-HA magnetic beads (ThermoFisher). The beads were washed three times with lysis buffer to reduce nonspecific binding. Bound proteins were eluted using HA peptide competition, and the eluates were concentrated using Amicon® Ultra-0.5 mL centrifugal filter units (Millipore). The concentrated NAMPT proteins were subsequently used in NAMPT enzymatic activity assays.

### Generation of SUDHL-1 and SUPM2 NAMPT-HA-GFP Stable Cell Lines

To generate stable cell lines expressing wild-type or mutant forms of NAMPT, SUDHL-1 and SUPM2 cells were transduced with lentiviral particles encoding NAMPT-HA-GFP (wild-type), NAMPT-Y188F-GFP, or the NAMPT-SS→AA-GFP dimerization mutant, as described above. Transductions were performed by mixing the ready-to-use lentiviral supernatant with 50,000 cells per well in a 96-well plate in the presence of 8 μg/mL polybrene (Millipore Sigma, catalog #TR-1003-G) to enhance infection efficiency. After a 16-hour incubation at 37°C, the media was replaced with fresh RPMI 1640 supplemented with 10% FBS, and the cells were cultured under standard conditions. Stable integration and expression of the constructs were monitored by GFP fluorescence, and GFP-positive cells were sorted and enriched by flow cytometry several days post-transduction to ensure homogeneity of expression.

### Plasmids and Lentiviral Reagents

Plasmids used in this study were obtained from Addgene: pHAGE-ALK (Addgene plasmid #116712); pHAGE-ALK-R1275Q (Addgene plasmid #116111); HIR WT (Human Insulin Receptor wild-type, Addgene plasmid #24049); pcDNA3.1 IGF1R-BirA(R118G)-HA (Addgene plasmid #109232). For lentivirus production, the Packaging Mix & Lentifectin Combo Pack was purchased from Applied Biological Materials (ABM, catalog #LV053-G074). Viral particles were concentrated using the Lenti-X™ Concentrator (Takara, catalog #631231) according to the manufacturer’s instructions.

### ALCL cell lines

SUPM2, SR786 and SUDH-L1 were purchased from the ATCC. The human CD4+cells transduced with NPM1::ALK, NA1 was created by our group as described earlier(*69*). The cell lines were regularly tested for Mycoplasma contamination using Mycoplasma detection kits from Thermo Fisher Scientific. All cells were grown in RPMI medium supplemented with 10% FBS and 1% penicillin/streptomycin under a humidified 37°C/5% CO_2_ incubator.

### ALK inhibitor (ALKi) resistant ALCL cell lines

ALKi crizotinib-resistant cell lines described earlier(*70*), KARPAS299 CRO6 (resistant up to 600nM of crizotinib and cultured at that concentration), SUPM2 CR03 (resistant up to 300nM of crizotinib and cultured at that concentration), and ALKi lorlatinib-resistant cell lines described earlier(*71*), KARPAS LR1000 (resistant up to 1000nM of lorlatinib and cultured at 300nM of lorlatinib) and SUPM2 LR1000 ((resistant up to 1000nM of lorlatinib and cultured at 300nM of lorlatinib). All cells these cell lines were obtained from Drs. Carlo Gambacorti-Passerini and Luca Mologni, University of Milano-Bicocca, Italy. The cell lines were grown in RPMI medium supplemented with 10% FBS and 1% penicillin/streptomycin under a humidified 37°C/5% CO_2_ incubator.

### Immunohistochemistry (IHC) of ALK+ALCL patient-derived biopsy samples

Human tissue microarrays from patients with ALCL (n = 8) were used to examine phospho-NAMPT Y188 and NAMPT. Expression was analyzed according to the samples’ pathology grade. Tissues were collected, fixed in 10% phosphate-buffered formaldehyde (formalin) for 24-48 hrs, dehydrated, and embedded in paraffin. Hematoxylin and eosin (H&E)-stained sections were used for morphological evaluation, and unstained sections were used for IHC studies. IHC staining was performed on a VENTANA Discovery XT automated staining instrument (Ventana Medical Systems) using VENTANA reagents according to the manufacturer’s instructions. Slides were deparaffinized using EZ Prep solution (cat # 950–102) for 16 min at 72 °C. Epitope retrieval was accomplished with CC1 solution (cat # 950–224) at high temperature (e.g., 95–100 °C) for 32 min. Rabbit primary antibodies (phospho-NAMPT Y188: 1:200, custom-developed by us; NAMPT, ProteinTech, Cat. 17183-AP) were titrated with a TBS antibody diluent and loaded into user-fillable dispensers for use on the automated stainer. The immune complex was detected using the Ventana OmniMap anti-rabbit detection kit (760–4311) and developed using the VENTANA ChromMap DAB detection kit (cat # 760-159) according to the manufacturer’s instructions. Slides were then counterstained with hematoxylin II (cat # 790-2208) for 8 min, followed by Bluing reagent (cat # 760-2037) for 4 min. Slides were then dehydrated through an ethanol series, cleared in xylene, and mounted. As a negative control, the primary antibody was replaced with normal rabbit IgG to confirm the absence of specific staining. Stained slides were scanned using an Aperio ScanScope CS 5 slide scanner (Aperio, Vista, CA, USA). Scanned images were then viewed and captured with Aperio’s image viewer software (ImageScope, version 11.1.2.760, Aperio).

### NAMPT enzymatic activity assay

NAMPT enzymatic activity was measured using the CycLex® NAMPT Colorimetric Assay Kit Ver.2 (MBL International; catalog no. CY-1251V2) according to the manufacturer’s instructions. HA-tagged NAMPT-WT and NAMPT-Y188F were expressed in HEK293T cells and ALK+ ALCL cell lines, SUDHL-1 and SUPM2 and immunopurified using anti-HA affinity purification. Purified NAMPT proteins were subjected to enzymatic activity assays using the supplied assay reagents, and NAMPT activity was quantified by measuring colorimetric signal generation according to the manufacturer’s protocol. NAMPT enzymatic activity was normalized to the amount of immunopurified NAMPT protein. Data represents at least three independent experiments.

### Quantitative reverse transcription PCR (qRT–PCR)

RNA extraction and quantitative reverse transcription PCR (qRT–PCR) were performed as previously described(*72*). Briefly, parental SUP-M2 and Karpas299 cells, as well as crizotinib- and lorlatinib-resistant derivatives (CRO3 and LR1000), were cultured in RPMI medium. Parental cells were maintained under standard culture conditions, whereas resistant cell lines were maintained in RPMI supplemented with 300 nM crizotinib or 200 nM lorlatinib, respectively. Cells were seeded in six-well plates and cultured for 48 h prior to RNA isolation.Total RNA was extracted and treated with DNase using the RNeasy Mini Kit and RNase-Free DNase Set (Qiagen; catalog nos. 74104 and 79254) according to the manufacturer’s instructions. RNA concentration was determined, and 1 µg of total RNA was used for cDNA synthesis using SuperScript IV Reverse Transcriptase (Invitrogen; catalog no. 18091050). qRT–PCR was performed using 1 µl of cDNA template and SYBR Green PCR Master Mix (Applied Biosystems; catalog no. A25742). Each 10 µl reaction contained 5 µl of SYBR Green mix, 0.5 µl of forward primer (10 mM stock), 0.5 µl of reverse primer (10 mM stock), 1 µl of cDNA, and 3 µl of nuclease-free water. Primer sequences are listed below. **NAMPT gene**: NAMPT-F: CGG-CAG-AAG-CCG-AGT-TCA-A; NAMPT-R: GCT-TGT-GTT-GGG-TGG-ATA-TTG-TT. **SLC12A8 gene**: SLC12A8-F: CTG-GTG-TCC-TTC-GTC-ATC-CTG; SLC12A8-R: CAC-CTG-CAA-CAC-ACT-GTC-CA. **SLC25A51 gene**: SLC25A51-F: AGGTCCTCTTTCGACAACAGC; SLC25A51-R: ACCAAACATAAGTGCAAGCGTA.

### Multiplex Immunofluorescence Staining and Image Acquisition

Multiplex immunofluorescence (mIF) staining was performed on formalin-fixed, paraffin-embedded (FFPE) tissue sections using the Discovery Ultra automated staining platform (Roche Diagnostics, Tucson, AZ, USA). Tissue sections were deparaffinized using EZ Prep solution (Roche Diagnostics; Cat. No. 950-102), followed by heat-induced epitope retrieval with Cell Conditioning 1 (Discovery CC1; Roche Diagnostics; Cat. No. 950-500) for 64 minutes. Sequential staining was performed using three primary antibodies and the OmniMap anti-Rabbit HRP detection system (Roche Diagnostics; Cat. No. 760-4311). ALK was detected using a ready-to-use rabbit monoclonal antibody (Roche Diagnostics; Cat. No. 790-4796) with incubation at 37°C for 60 minutes. NAMPT was detected using a rabbit polyclonal antibody (Proteintech; Cat. No. 11776-1-AP) at a dilution of 1:5,000 and incubated at room temperature for 32 minutes. VDAC1 was detected using a rabbit polyclonal antibody (Proteintech; Cat. No. 55259-1-AP) at a dilution of 1:1,500 and incubated at room temperature for 32 minutes. Antibody denaturation steps were performed between staining cycles according to the manufacturer’s multiplex staining protocol to minimize cross-reactivity and allow sequential detection of multiple rabbit primary antibodies. Target visualization was achieved using the DISCOVERY FAM Kit (Roche Diagnostics; Cat. No. 760-243), DISCOVERY Rhodamine 6G (R-6G) Kit (Roche Diagnostics; Cat. No. 760-244), and DISCOVERY Cy5 Kit (Roche Diagnostics; Cat. No. 760-238) for ALK, NAMPT, and VDAC1, respectively. Nuclei were counterstained with DAPI. Following staining, slides were coverslipped using a fluorescence-compatible mounting medium and scanned using a NanoZoomer S60 digital slide scanner (Hamamatsu Photonics, Hamamatsu, Japan). Whole-slide fluorescent images were acquired at 40× magnification using DAPI, FAM, R-6G, and Cy5 filter sets. Digital images were reviewed for staining quality and subsequently analyzed using digital image analysis software.

### Native NuPAGE Gel Electrophoresis to Assess NAMPT Dimerization

To assess the dimerization status of NAMPT, native gel electrophoresis was performed using the NativePAGE™ Bis-Tris Gel System (Thermo Fisher Scientific), which enables the resolution of protein complexes under non-denaturing conditions. SUPM2 ALK⁺ ALCL cells expressing HA-tagged NAMPT constructs including wild-type (NAMPT-WT-HA), a phosphorylation-deficient mutant (NAMPT-Y188F-HA), and a dimerization-defective mutant (NAMPT-SS-AA-HA) were lysed on ice in NativePAGE™ Sample Buffer supplemented with 0.5% digitonin or 0.1% Triton X-100, along with protease and phosphatase inhibitors, for 30 minutes with gentle rotation according to the instructions manuel. Lysates were clarified by centrifugation at 14,000 × g for 15 minutes at 4 °C. Protein concentrations were determined using a BCA assay, and equal amounts of total protein (20–30 µg) were mixed with NativePAGE™ 4× Sample Buffer and G-250 additive according to the manufacturer’s instructions. Samples were resolved on a NativePAGE™ 4–16% Bis-Tris gel using NativePAGE™ Running Buffer. The initial run was conducted with Dark Blue Cathode Buffer at 150 V for 30–45 minutes, followed by Light Blue Cathode Buffer at 200 V for ∼90 minutes at 4 °C. After electrophoresis, proteins were transferred onto a nitrocellulose membrane using NuPAGE™ Transfer Buffer containing 10% methanol, at 90 V at 4 °C. Membranes were blocked in 5% non-fat milk in TBST and probed with anti-HA antibody to detect total NAMPT, and a phospho-specific NAMPT-Y188 antibody to assess phosphorylation in the context of dimerization. Protein bands were visualized using HRP-conjugated secondary antibodies and chemiluminescence on an iBright Western blot imaging system. As a positive control for native protein complex migration, HA-tagged ACLY, a known tetrameric enzyme, was included. Native molecular weight standards (NativeMark™, Thermo Fisher) were also loaded to estimate the molecular weight of monomeric and dimeric NAMPT forms.

### *In-vitro* kinase assay and detection of NAMPT tyrosine phosphorylation

Recombinant human NAMPT protein and a panel of active recombinant human tyrosine kinases (TKs) were used to assess NAMPT phosphorylation in vitro. The tyrosine kinases included ALK, SRC, FYN, LCK, BTK, FLT3, ABL, SYK, and EGFR. In addition, non-tyrosine kinases AKT1 and ERK1 were included as negative controls for comparison. In vitro kinase assays were performed by incubating recombinant NAMPT with each kinase in kinase reaction buffer under conditions optimized for each enzyme. Reactions were terminated by adding SDS loading buffer, followed by SDS-PAGE and immunoblotting. Tyrosine phosphorylation of NAMPT was detected using a phosphotyrosine-specific antibody (pY100), enabling assessment of kinase-specific phosphorylation patterns.

### Large-scale in vitro kinase assay and phosphoproteomic analysis of NAMPT

To identify specific tyrosine phosphorylation sites on NAMPT, we performed large-scale in vitro kinase assays using two active tyrosine kinases, SYK and BTK, selected based on preliminary screening. Recombinant human NAMPT was incubated with either SYK or BTK under optimized kinase reaction conditions, as described above. Following the reaction, samples were run on a NuPAGE 10 % Tris-glycine gel for ∼0.5–1 cm migration from the well, until proteins enter the stacking gel, followed by colloidal Coomassie staining. NAMPT bands were excised for in-gel digestion and phosphopeptide enrichment. Phosphorylation sites were identified by liquid chromatography-tandem mass spectrometry (LC-MS/MS), as previously described(*73*).

To investigate the impact of tyrosine phosphorylation and dimerization on NAMPT’s interactome in a cellular context, we performed proteomic analyses using liquid chromatography–tandem mass spectrometry (LC–MS/MS). To assess the role of dimerization, we generated a dimerization-defective NAMPT mutant by substituting alanine for serine residues at positions 199 and 200 (S199A/S200A), thereby disrupting the dimer interface. HA-tagged wild-type (NAMPT-WT-HA) and mutant (NAMPT_SS-AA-HA) constructs were introduced into HEK293T cells via lentiviral transduction to generate stable cell lines. These were then transfected with either an empty vector (N=1), NAMPT-WT-HA alone (N=3), or in combination with constitutively active NPM–ALK (N=3). Similarly, NAMPT_SS-AA-HA was expressed alone (N=3) or co-expressed with NPM–ALK (N=3). Following the expression, NAMPT-containing protein complexes were immunoprecipitated using anti-HA magnetic beads.

Mass spectrometry of Immunoprecipitated samples were run 0.5 cm into a 10% SDS-gel and subjected to in-gel trypsin digestion. Tryptic digests were resuspended in 0.015% dodecyl maltoside, 0.1% trifluoroacetic acid before analysis. LC-MS/MS analysis was performed using an Orbitrap Astral mass spectrometer (Thermo Fisher Scientific) coupled to a Vanquish Neo UPLC system (Thermo Fisher Scientific). Chromatographic separation was carried out at 45.0 °C with a flow rate of 300 nL/min. Buffer A consisted of 0.1% formic acid in Milli-Q water, and buffer B consisted of 0.1% formic acid in acetonitrile. Peptides were loaded onto a trap column (100 Å, 75 μm i.d. x 2 cm packed with 3 μm C18 resin; Thermo Fisher Scientific), which was connected in line with a nanoEase M/Z Peptide BEH C18 nanocapillary analytical column (130 Å, 75 μm i.d. x 25 cm, 1.7 μm particle size; Waters). Separation was achieved using a 40 min linear gradient: 5-30% B over 25 min, 30-40% B over 6 min, 40-80% B over 2 min, followed by a 7 min hold at 80% B before re-equilibration. MS data were acquired in data-independent acquisition mode. An MS1 scan was collected every 0.6 s in the Orbitrap at 240,000 resolution. Ions were injected for 3 ms or until an AGC target of 5e6 was reached. Precursor ions within a mass range of 380-980 *m/z* were collected in 2 *m/z* isolation windows with a maximum injection time of 3 ms or until an AGC target of 5e4 ions was reached. Precursors were fragmented using 25% normalized collisional energy. MS2 scans from 110 – 2000 *m/z* were collected in the Astral mass analyzer.

### Data Analysis

Mass spectrometry data were processed using DIA-NN v2.1.0 and searched against a spectral library generated from the SwissProt human database, supplemented with the recombinant NAMPT sequence and a common contaminants database(*74*). N-terminal acetylation, N-terminal methionine excision, S/T/Y phosphorylation, and methionine oxidation were set as variable modifications. Cysteine carbamidomethylation was set as a static modification Data were searched with full tryptic specificity allowing for 2 missed cleavages and 3 variable modifications, a peptide length of 7-50 amino acids, a precursor charge of 1-4, and a scan window of 15. MS1 and MS2 mass accuracy were set to 5.0 and 10.0 ppm, respectively. The match-between-runs feature was enabled, and the cross-run normalization feature was disabled.

Contaminants, proteins identified by a single peptide, and proteins with fewer than 3 valid values in any of the conditions were removed from the filtered dataset. Protein abundance values were normalized to the median total abundance using Normalyzer DE(*75*). Differential expression analysis was performed using LFQ Analyst(*76*). Missing values were imputed with the minimum value of the filtered dataset, and FDR levels were controlled using the Benjamini-Hochberg correction. Proteins with an absolute fold change > 2 and an adjusted p-value < 0.05 were considered significant. Pathway enrichment analysis including gene ontology analysis, KEGG pathway analysis, and STRING network analysis of significantly enriched proteins analysis was performed using Pathway-Analyst (add reference PMID: 31657565). Site-level phosphorylation identifications used a 90% confidence localization cutoff.

### Metabolomics analysis

Metabolomic profiling was performed for ALK⁺ ALCL cell lines treated with either DMSO or the ALK inhibitor lorlatinib. Metabolite extraction and analysis were carried out at The Wistar Institute Proteomics and Metabolomics Shared Resource. Extractions were performed for 15 million cells per sample using 10:3:2.5 (v/v/v) MTBE/methanol/water spiked with an isotopically-labeled metabolite standard mix (MSK-A2-1.2; Cambridge Isotope Laboratories). The lower aqueous phase was dried, resuspended in 80% methanol, and stored at -80 °C. LC-MS analysis was performed as described previously with modifications(*77*). Briefly, samples were analyzed by LC-MS/MS on a Thermo Scientific Q Exactive Plus mass spectrometer with in-line Vanquish UHPLC System. LC separation was performed using a ZIC-pHILIC column, 150 × 2.1 mm, 5 μM, maintained at 45 °C (EMD Millipore). Mobile phase A was 20 mM ammonium carbonate, 5 µM medronic acid, 0.1% ammonium hydroxide, pH 9.2, and mobile phase B was acetonitrile. Analytical separation was performed at 0.2 ml/min flow rate using the following gradient: 0 min, 85% B; 2 min, 85% B; 17 min, 20% B; 17.1 min, 85% B; and 26 min, 85% B. Samples were analyzed by full MS scans with polarity switching (for all samples) or full MS/data-dependent MS/MS scans with separate acquisitions for positive and negative polarities (for sample pool, QC). Relevant MS parameters include: sheath gas, 30; auxiliary gas, 5; sweep gas, 0; auxiliary gas heater temperature, 200 °C; spray voltage, 3.6 kV; capillary temperature, 325 °C; S-lens RF, 65. Full MS scans were acquired using a scan range of 65 to 975 m/z; 70,000 resolution; automated gain control (AGC) target of 1E6; and maximum injection time (IT) of 100 ms. Data-dependent MS/MS was performed on the 10 most abundant ions; 17,500 resolution; AGC target of 5E4, maximum IT of 50 ms, isolation width of 1.0 m/z, and stepped normalized collision energy of 20, 40, 60.

Raw data were processed using Compound Discoverer 3.4 (Thermo Scientific) with separate analyses for positive and negative polarities. Metabolites were identified by matching accurate mass and retention time to standards or by querying MS/MS scans against the mzCloud spectral database (full match, score > 50; https://mzCloud.org). Metabolite levels were quantified relatively by integrated peak area using full MS data. Metabolite levels were corrected for instrument drift using peak areas from technical injections of the QC pool run throughout the sample set and to the total MS signal from identified metabolites in each sample.

### NAD⁺ quantification assay

Intracellular NAD⁺ levels were measured using the NAD/NADH Assay Kit (Abcam; catalog no: ab65348) according to the manufacturer’s instructions. ALK inhibitor–sensitive (ALKi-S) and ALK inhibitor–resistant (ALKi-R) SUP-M2 and Karpas299 cells were treated with lorlatinib, FK866, or the combination for 48 h. Cells were harvested, lysed in the supplied extraction buffer, and total NAD⁺ was quantified using the enzymatic cycling assay. NAD⁺ concentrations were calculated from a standard curve and normalized to total protein content, determined by BCA assay, and expressed as ng NAD⁺/mg protein. Data represents at least three independent biological experiments.

### Cell proliferation assay

Cell proliferation was assessed using the WST-1 Cell Proliferation Assay ((Millepore-sigma, catalog# 5015944001) according to the manufacturer’s instructions. ALK inhibitor–sensitive (ALKi-S) and ALK inhibitor–resistant (ALKi-R) SUP-M2 and Karpas299 cells were seeded at equal densities in 96-well plates and treated with lorlatinib, FK866, or the combination for 72 h. WST-1 reagent was then added directly to each well and incubated according to the manufacturer’s protocol. Absorbance was measured using a microplate reader at 450 nm. Cell proliferation was expressed relative to vehicle-treated controls. Data represents at least three independent biological experiments performed with technical replicates.

### Molecular Docking of NAMPT and NPM–ALK in the Presence of Nicotinamide

To investigate the potential structural interface between NAMPT and the oncogenic tyrosine kinase NPM–ALK, molecular docking simulations were performed in the presence of the NAMPT substrate nicotinamide (NAM). The goal was to model how substrate binding alters the conformational landscape of NAMPT to facilitate interaction with NPM–ALK. The crystal structure of human NAMPT in complex with nicotinamide (PDB ID: *e.g., 2GVJ*) was used as the receptor. The NPM–ALK fusion protein structure was modelled using available ALK kinase domain coordinates (PDB ID: *e.g., 3LCS*) and the known NPM1::ALK fusion sequence, followed by energy minimization. Protein–protein docking was performed using HADDOCK 2.4 or ClusPro, incorporating substrate-bound NAMPT as the receptor. Key interaction residues were guided by published phosphorylation data and structural constraints. Docking scores, interface residues, and predicted binding energies were analyzed to evaluate the impact of nicotinamide binding on NAMPT’s ability to engage NPM–ALK. The docking results suggested that nicotinamide-bound NAMPT adopts a conformation that exposes potential interaction surfaces, favoring stable complex formation with the kinase domain of NPM–ALK. This supports the hypothesis that substrate occupancy may prime NAMPT for phosphorylation by oncogenic tyrosine kinases.

### Statistics and reproducibility

The student’s *t*-test was used to analyze differences in western blot densitometric values and in the activity assay to assess differences in cell growth and colony formation. *p-*values equal to or less than 0.05 were considered statistically significant without being adjusted for multiple comparisons.

## Data availability

We confirm that all relevant data and methods are included in the main article, and the supplementary material of proteomics will be uploaded to the appropriate website. The mass spectrometry data generated in this study have been deposited in the ProteomeXchange Consortium through the MassIVE repository under the dataset identifier PXD066479.Reviewer login credentials:FTP URL: ftp://MSV000098617@massive-ftp.ucsd.edu USN: MSV000098617_reviewer PW: NAMPT

## Acknowledgments

We thank the Proteomics and Metabolomics Shared Resource at the Wistar Institute, Philadelphia, for providing metabolomics and proteomics analysis services and for their support. We thank Jirong Zhang for assistance with IHC staining. We thank Lori Rink, lab members Jane Koshy, Delia Zumpano, and Adam Chatoff for providing cell lysates and TMA slides. The figures in this manuscript were created using the Biorender program. H.Y.T is supported by R50CA221838. Funding support for The Wistar Institute’s Proteomics and Metabolomics Shared Resource was provided by Cancer Center Support Grant P30 CA010815. No NIH or other US federal funds were used to support co-authors with foreign affiliations.

## Author Contributions

J.B. conceived the project, designed and performed the research, analyzed the data, supervised the project, and wrote the manuscript. C.L., C.A., D.R., and N.S. performed the experiments. A.M.F. and H.Y.T. conducted the proteomics analysis. L.W., K.Q.C., and R.N contributed to the IHC and IF. A.E. performed confocal imaging. A.R.G. and W.Z conducted metabolomics. J.L.S., L. Rink, L.M., A.N.H., J.A.B., and J.C. contributed key reagents. R.D. performed molecular docking. M.W. obtained funding, patient samples, provided resources, analyzed the data, and revised the manuscript. All authors reviewed and approved the manuscript.

## Supplementary Figures

**Fig. S1.**
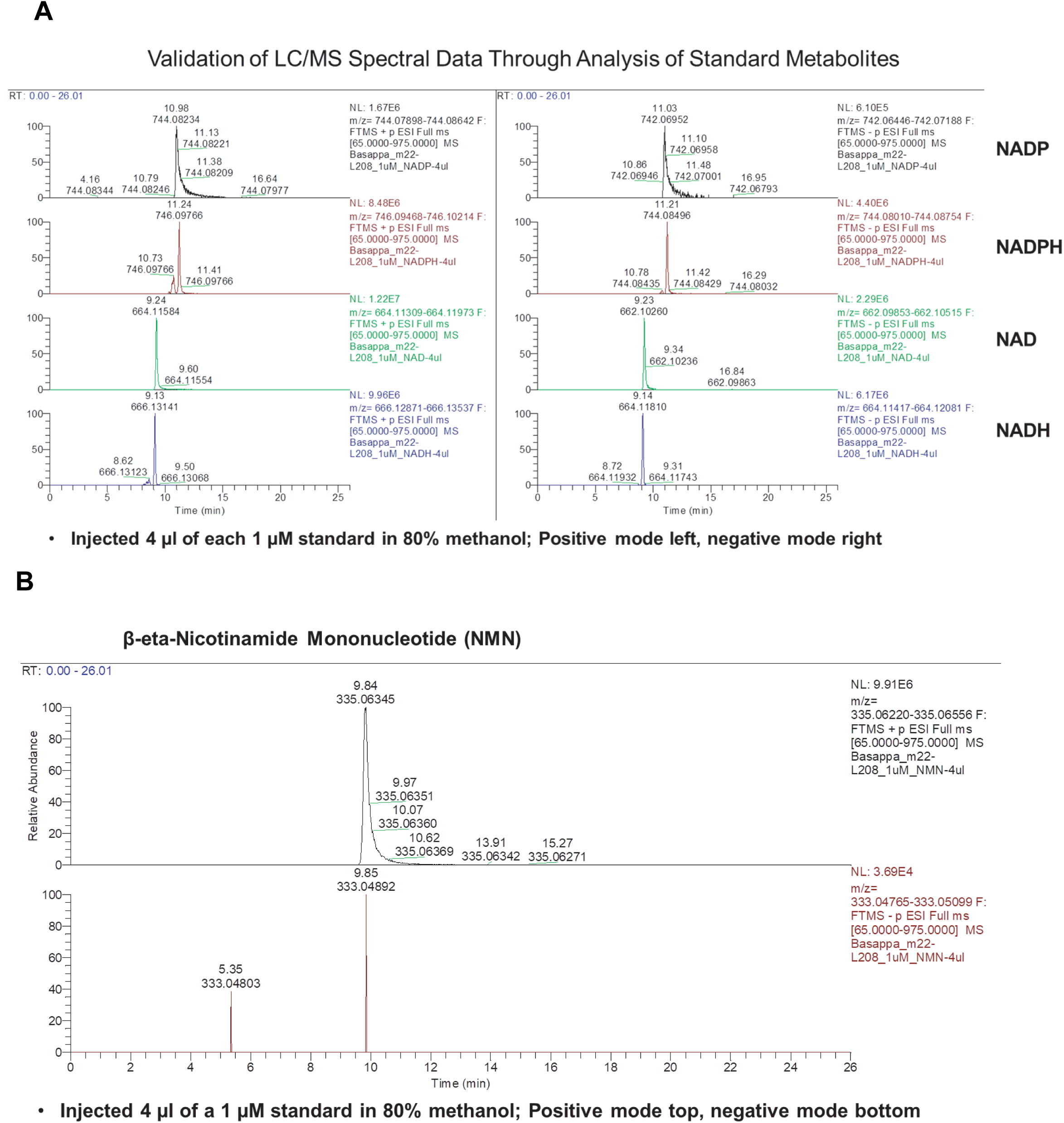
Validation of LC–MS/MS metabolite using standard metabolites. **(A and B)** Representative chromatograms and calibration analyses generated using authentic standards for NAD⁺, NADH, NADP⁺, NADPH, and NMN, confirming metabolite identity and quantification accuracy.

**Fig. S2.**
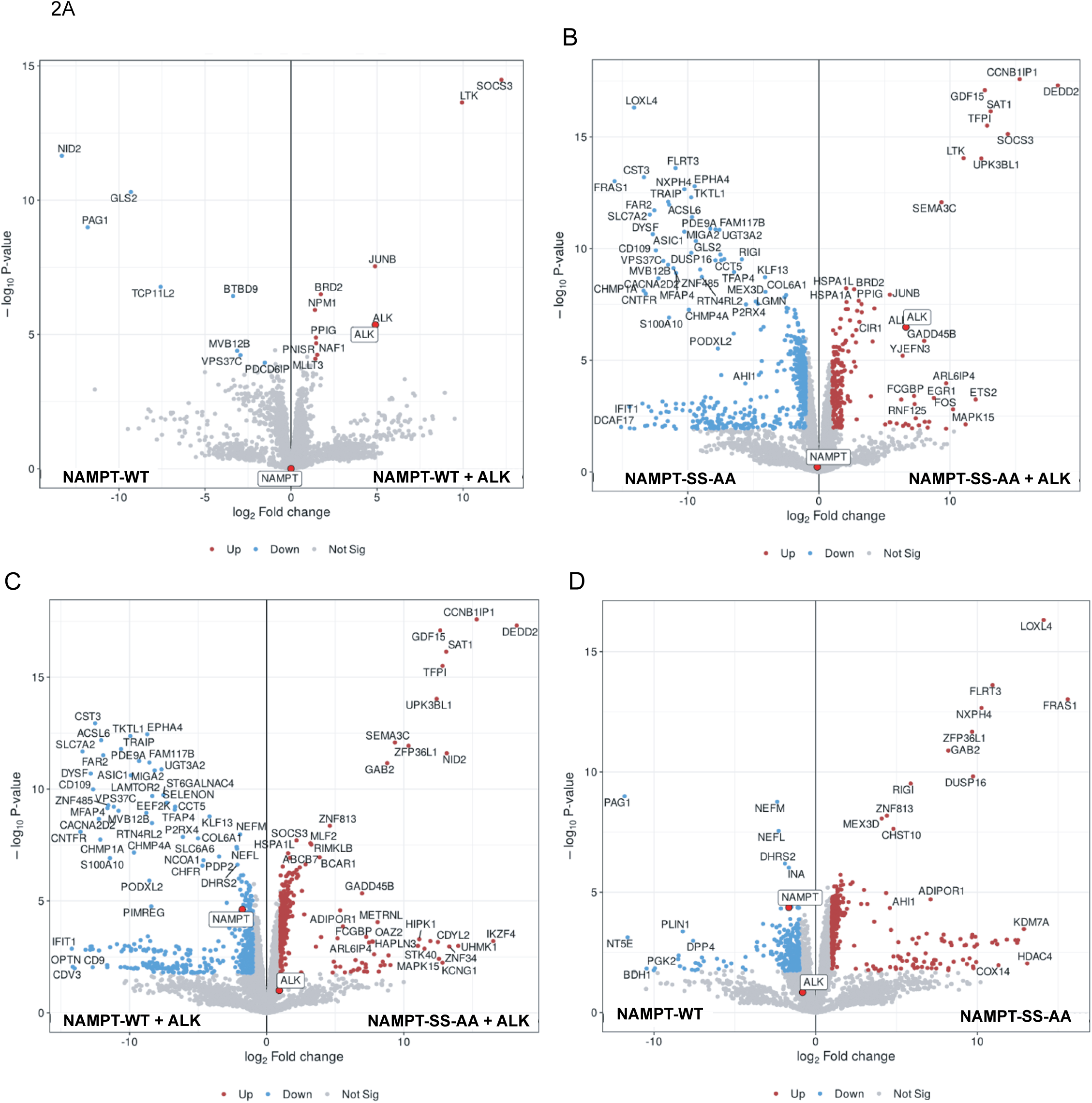
Comparative analysis of the NAMPT interactome across phosphorylation and ALK-expression states. **(A to D),** Volcano plots showing differential enrichment of NAMPT-associated proteins across the indicated comparisons: (**A)** NAMPT-WT versus NAMPT-WT+ALK; (**B)** NAMPT-SS-AA versus NAMPT-SS-AA+ALK; (**C)** NAMPT-WT+ALK versus NAMPT-SS-AA+ALK; and (**D)** NAMPT-WT versus NAMPT-SS-AA. Each point represents an identified interactor quantified by mass spectrometry. The x-axis indicates log2 fold change, and the y-axis indicates −log10 (*P*) value. Proteins significantly enriched in each condition are highlighted. These analyses reveal phosphorylation- and ALK-dependent remodeling of the NAMPT interactome, identifying distinct subsets of proteins that preferentially associate with specific NAMPT phosphorylation states and ALK signaling contexts.

**Fig. S3.**
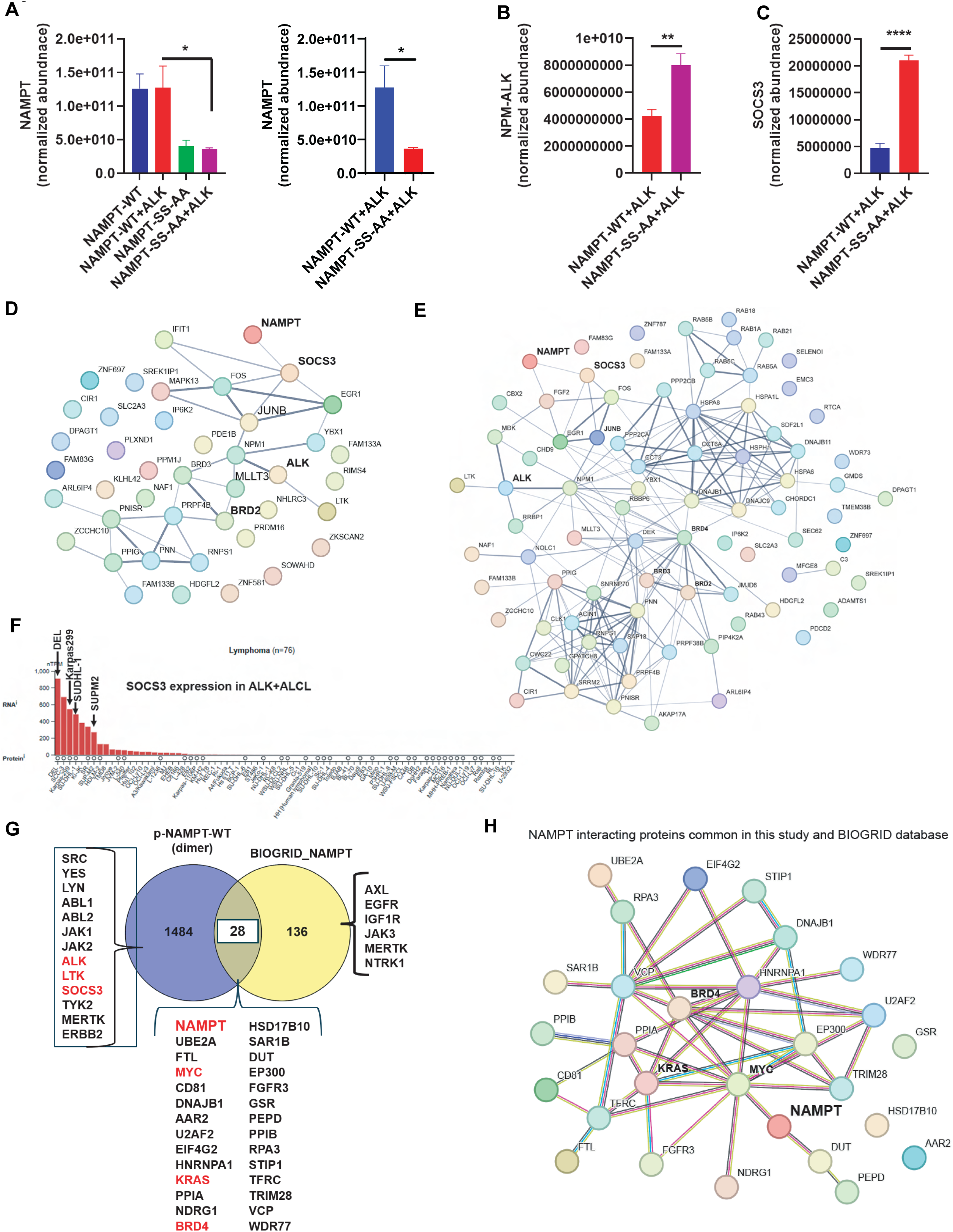
Phosphorylation-dependent remodeling of the NAMPT interactome. (**A)** Relative abundance of monomeric and dimeric NAMPT in immunoprecipitated bait protein and proteomic datasets, showing that dimeric NAMPT is substantially more abundant than monomeric NAMPT. **(B)** Despite its lower abundance, NPM–ALK preferentially associates with Y188-phosphorylated monomeric NAMPT, indicating a phosphorylation and conformation-dependent interaction independent of protein abundance. Y188 is located at the NAMPT dimer interface, and disruption of dimerization by the SS–AA mutant is predicted to increase Y188 accessibility, facilitating phosphotyrosine-dependent recognition by NPM–ALK. (**C)** The SH2 domain-containing adaptor SOCS3 similarly exhibits preferential binding to Y188-phosphorylated monomeric NAMPT, consistent with phosphotyrosine-dependent substrate recognition. (**D)** Proteomic analysis identifies the epigenetic reader BRD2 as a prominent interactor enriched in the Y188-phosphorylated NAMPT interactome relative to non-phosphorylated NAMPT. (**E and F)** STRING network analysis of proteins significantly enriched in the p-NAMPT Y188 interactome (log₂ fold change >2, *P* < 0.05) reveals a prominent ALK–NAMPT–SOCS3 signaling network, supporting the biochemical interactions identified by affinity proteomics. (**G)** Analysis of publicly available Human Protein Atlas datasets demonstrates high SOCS3 expression across ALK⁺ ALCL cell lines, consistent with activation of the ALK–SOCS3 signaling axis. (**H)** Comparison of the Y188-phosphorylated NAMPT interactome with the BioGRID database identifies 28 previously reported NAMPT-interacting proteins, including MYC, KRAS, BRD4, and BRD2, while revealing more than 1,484 previously unreported NAMPT-associated proteins. (**I)** STRING network analysis of NAMPT common interactome. (**J)** Co-immunoprecipitation and LC-MS/MS validates the interaction of BRD4 with both phosphorylated and non-phosphorylated NAMPT, indicating that BRD4 association is independent of Y188 phosphorylation status.

**Fig. S4.**
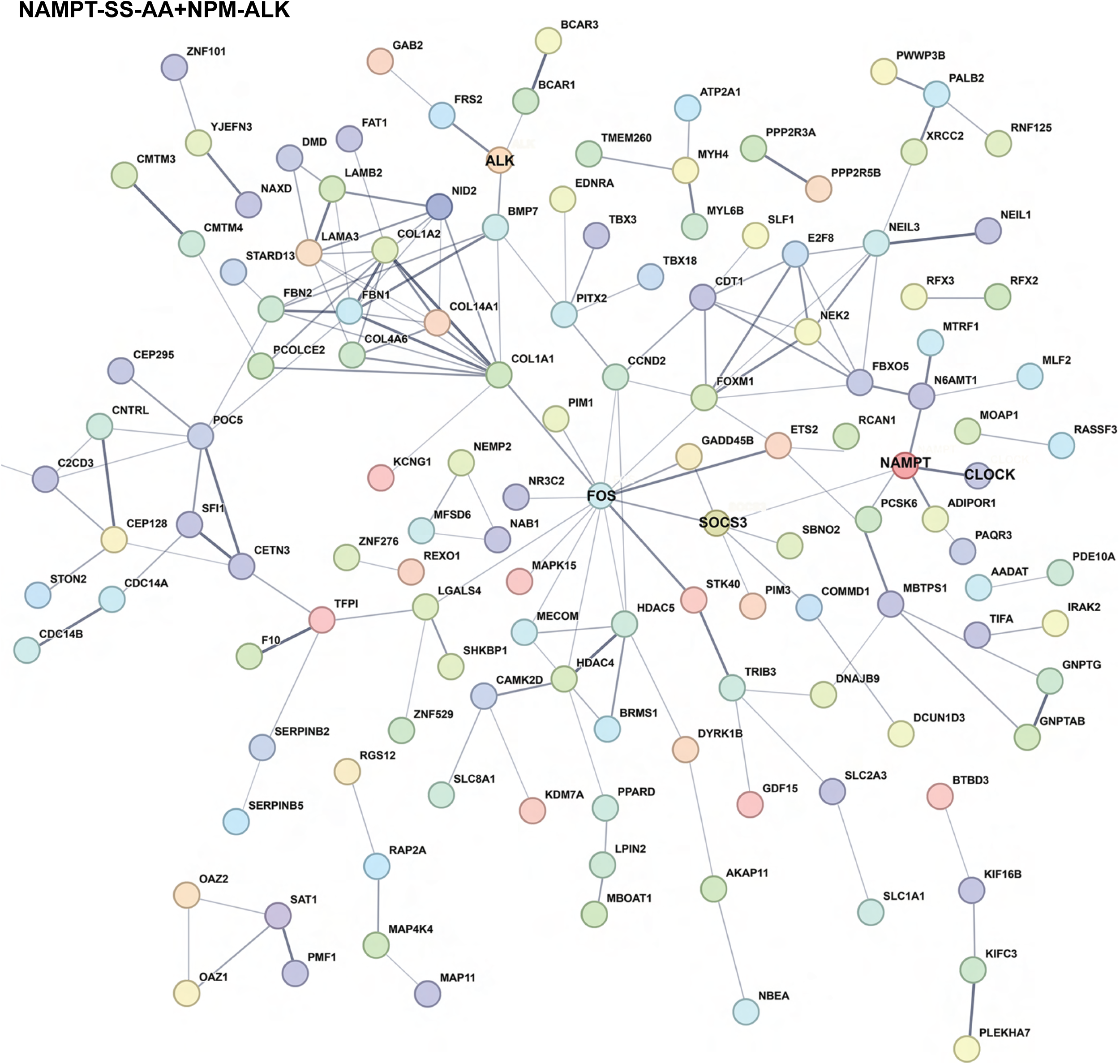
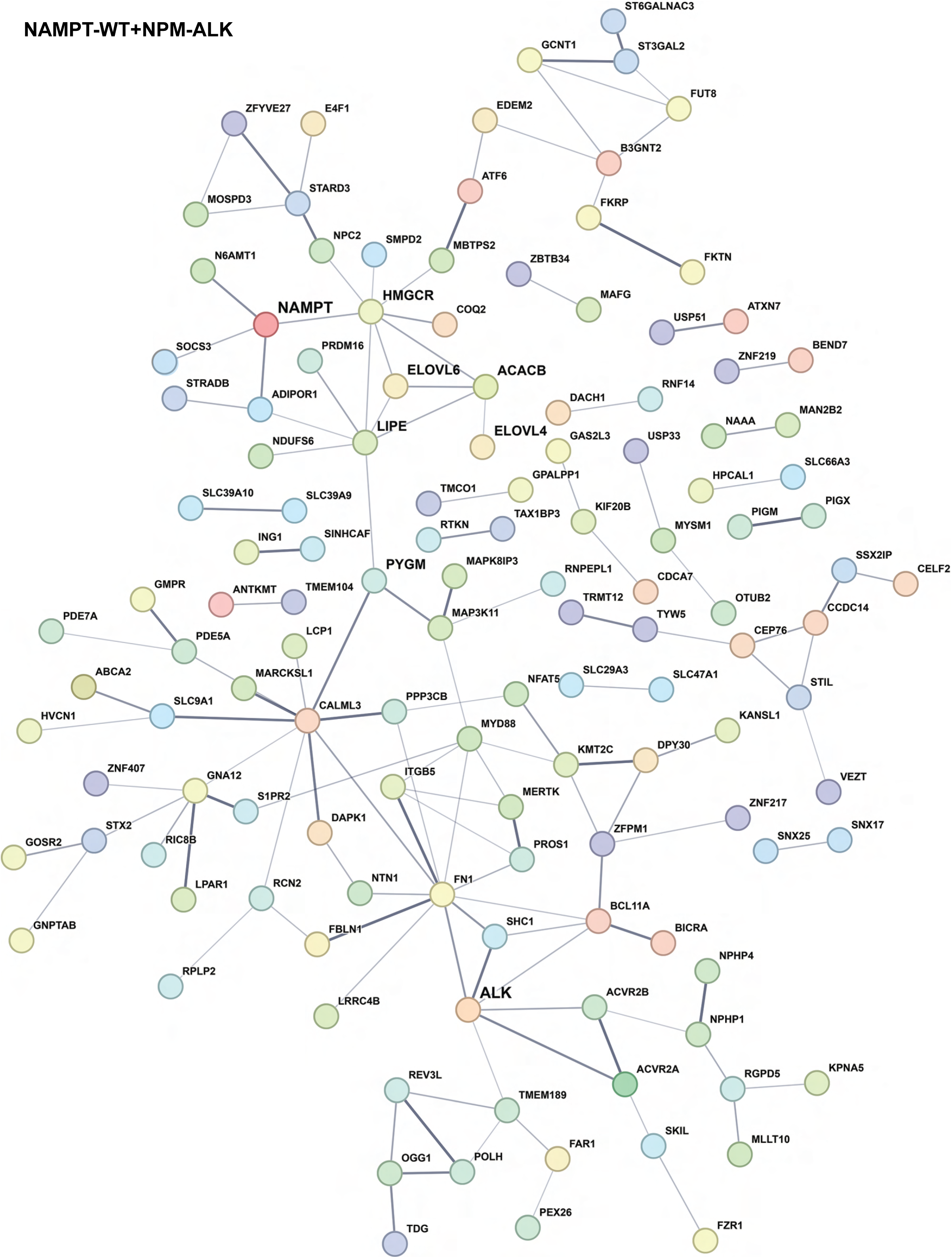
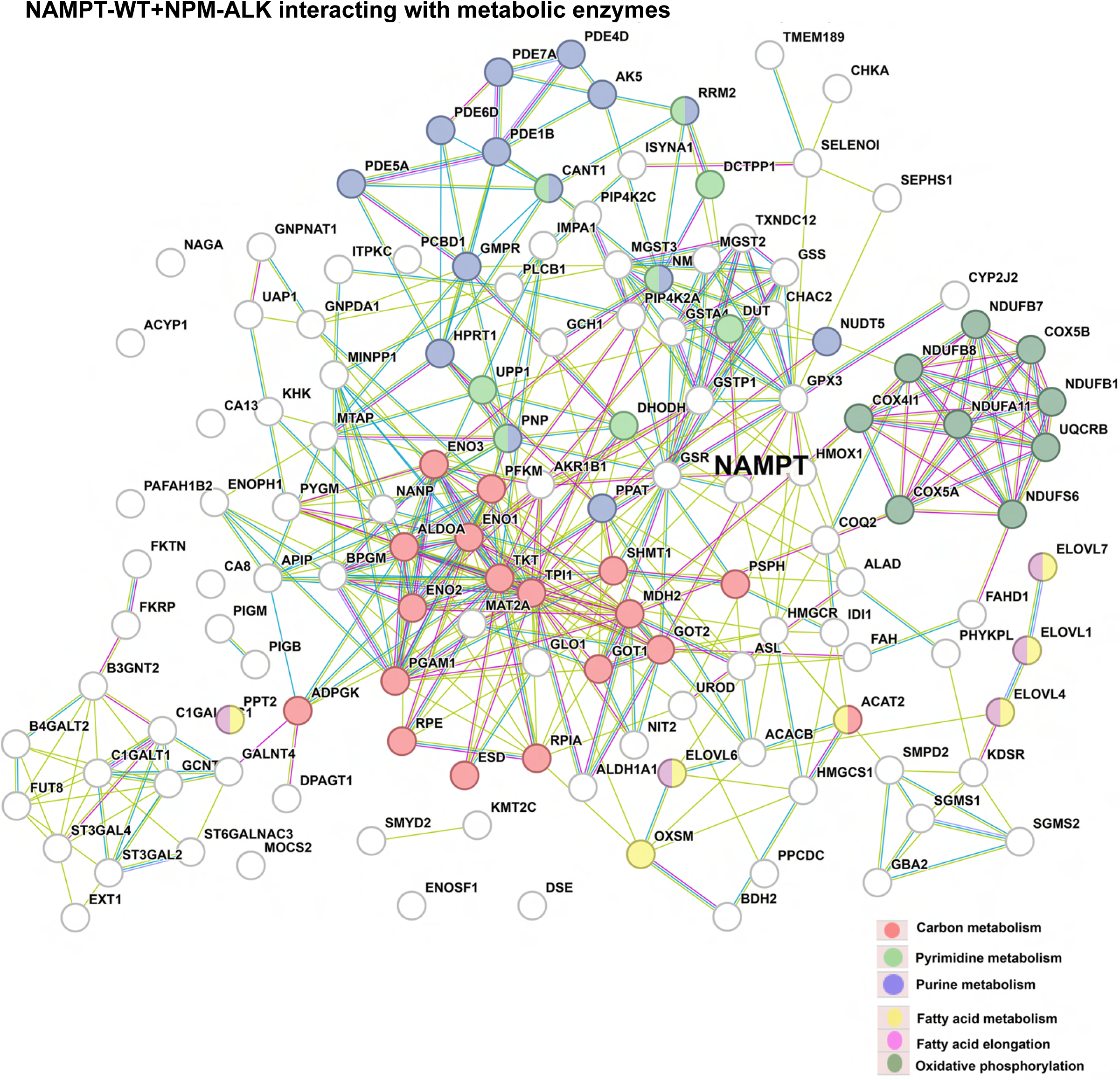
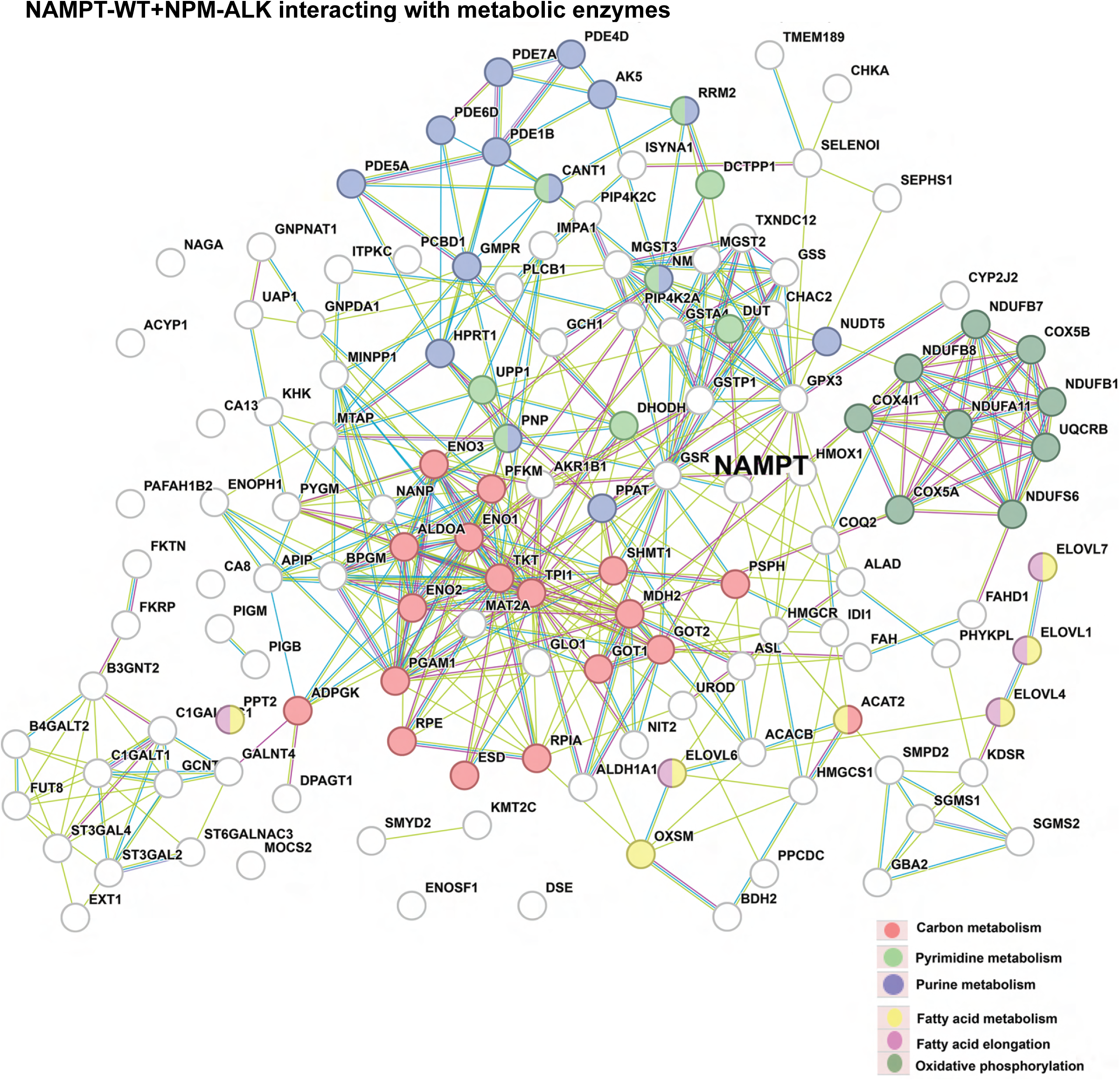
Phosphorylation-dependent NAMPT interactome. (**A)** STRING network analysis of proteins significantly enriched in the p-NAMPT Y188/NAMPT (monomer). **(B)** STRING network analysis of proteins significantly enriched in the p-NAMPT Y188/NAMPT (dimer). (**C)** STRING network analysis of proteins significantly enriched in the p-NAMPT Y188/NAMPT (dimer) by focusing on metabolic pathways enzymes. (**D)** STRING network analysis of proteins significantly enriched in the p-NAMPT Y188/NAMPT (dimer) phosphoproteins.

